# HSP90 modulates T2R bitter taste receptor nitric oxide production and innate immune responses in human airway epithelial cells and macrophages

**DOI:** 10.1101/2021.11.16.468387

**Authors:** Ryan M. Carey, Benjamin M. Hariri, Nithin D. Adappa, James N. Palmer, Robert J. Lee

## Abstract

Bitter taste receptors (T2Rs) are G protein-coupled receptors (GPCRs) expressed in various cell types including ciliated airway epithelial cells and macrophages. T2Rs in these two airway innate immune cell types are activated by bitter products, including some secreted by *Pseudomonas aeruginosa*, leading to Ca^2+^-dependent activation of endothelial nitric oxide (NO) synthase (eNOS). NO enhances mucociliary clearance and has direct antibacterial effects in ciliated epithelial cells and increases phagocytosis by macrophages. Using biochemistry and live cell imaging, we explored the role of heat shock protein 90 (HSP90) in regulating T2R-dependent NO pathways in primary sinonasal epithelial cells, primary monocyte-derived macrophages, and a human bronchiolar cell line (H441). We used immunofluorescence to show that H441 cells express eNOS and certain T2Rs and that the bitterant denatonium benzoate activates NO production in an HSP90-dependent manner in cells grown either as submerged cultures and at air liquid interface. In primary sinonasal epithelial cells, we determined that HSP-90 inhibition reduces T2R-stimulated NO production and ciliary beating which are crucial for pathogen clearance. In primary monocyte-derived macrophages, we found that HSP-90 is integral to T2R-stimulated NO production and phagocytosis of FITC-labeled *Escherichia coli* and pHrodo-*Staphylococcus aureus*. Our study demonstrates that HSP90 serves an innate immune role by regulating NO production downstream of T2R signaling by augmenting eNOS activation without impairing upstream calcium signaling. These findings suggest that HSP90 plays an important role in airway antibacterial innate immunity and may be an important target in airway diseases like chronic rhinosinusitis, asthma, or cystic fibrosis.

## INTRODUCTION

Bitter taste receptors (also known as taste family 2 receptors, or T2Rs, encoded by *TAS2R* genes) are G protein-coupled receptors (GPCRs) used by the tongue to detect bitter compounds (1, 2). However, many of the 25 human T2R isoforms are also expressed in other organs (1–5), including the nose, sinuses, and lungs (6, 7). These receptors regulate diverse processes like airway smooth muscle contraction (8–11) and innate immune responses in the oral epithelium (12). T2R receptors are also expressed in immune cells like monocytes and macrophages (13–15), which are important players in airway innate immunity (16, 17). In the airway epithelium, T2R isoforms 4, 14, 16, 38, and possibly others are expressed in bronchial and nasal motile cilia (18–21). These T2Rs are activated in response to acyl-homoserine lactone (AHL) and quinolone quorum-sensing molecules secreted by the common airway pathogen *Pseudomonas aeruginosa* (18, 22, 23).

Activation of the T2Rs in sinonasal cilia or unprimed (M0) macrophages causes Ca^2+^-dependent activation of nitric oxide (NO) synthase (NOS) (4, 18, 24–28), likely the endothelial NOS (eNOS) isoform expressed in both airway ciliated cells (29–32) and M0 macrophages (33). In ciliated cells, NO activates soluble guanylyl cyclase to produce cyclic GMP (cGMP). NO activates protein kinase G (PKG) to elevate ciliary frequency to increase mucociliary clearance, the major physical defense of the airway. The T2R-activated NO also directly diffuses into the airway surface liquid (ASL), where it can have antibacterial effects (18, 34). NO can damage bacterial cell walls and/or DNA (35, 36). NO can also inhibit the replication of many respiratory viruses, including influenza, parainfluenza, rhinovirus,(37) and SARS-CoV-1 & −2 (38–41). In macrophages, T2R-stimulated NO production and cGMP production acutely increases phagocytosis (42). Thus, T2R to NO signaling may also be an important therapeutic target in infectious diseases beyond the airway. As NOS and NO have been implicated in chronic rhinosinusitis (43) as well as asthma and other lung diseases (44), better understanding of the mechanisms of NOS activation in airway epithelial cells may have implications beyond T2Rs.

The importance of T2Rs in upper airway defense is supported by observations that patients homozygous for the AVI *TAS2R38* polymorphism, which renders the T2R38 receptor non-functional, have increased frequency of gram-negative bacterial infection (18), have higher levels of sinonasal bacteria in general (45, 46) and specifically have higher levels of biofilm-forming bacteria (47), have higher frequency of chronic rhinosinusitis (48–51), and exhibit worse outcomes after functional endoscopic sinus surgery (52). One study has suggested that *TAS2R38* genetics may also play a role in cystic fibrosis *P. aeruginosa* infection (53), though other studies have suggested that *TAS2R38* may not be a modifier gene in CF (54, 55). Recently, an association between the *TAS2R38* PAV (functional) genotype has been associated with a lower mortality of SARS-COV2 compared with the AVI (non-functional) genotype (56). Better understanding the role of T2R38 and other T2Rs in airway innate immunity is important for determining if and how to leverage these receptors as therapeutic targets or predictive biomarkers.

In this study, we explored the role of heat shock protein 90 (HSP90) in T2R function in two types of cells important for airway innate immunity: ciliated epithelial cells and macrophages. The members of the HSP90 class of molecular chaperones are highly conserved and ubiquitously expressed (57). In addition to promoting protein folding, HSP90 regulates signaling by facilitating trafficking or localization of signaling proteins and/or functioning as molecular scaffolds to bring signaling molecules together. Because HSP90 is critical for endothelial cell NO production via eNOS, we tested if HSP90 is involved in T2R-dependent NO generation. HSP90 proteins can facilitate eNOS activation via scaffolding of eNOS with activating kinases such as Akt or Ca^2+^-bound calmodulin (CaM) kinases (58–62). HSP90 has been localized to the base of airway cell cilia (63–65), suggesting HSP90 may be localized close to T2Rs in airway ciliated cells and may help facilitate their signal transduction to eNOS. Recently, it was proposed that HSP90 inhibition by geldanamycin can revert Th2- and Th17-induced airway epithelial goblet cell metaplasia (66). If HSP90 inhibition reduces T2R NO responses, this may have unwanted effects of reducing T2R/NO-mediated innate immunity.

To test the requirement for HSP90 in T2R signaling, we combined biochemistry and live cell imaging with a human bronchiolar cell line, primary sinonasal epithelial cells, and primary monocyte-derived macrophages. Results below reveal important molecular insights into the T2R signaling pathway in both airway epithelial and immune cells and identify a specific role for HSP90 in airway epithelial NO-mediated innate immunity.

## METHODS

### Cell culture

HEK293T cells were obtained from ATCC (ATCC Cat# CRL-3216, RRID:CVCL_0063) and cultured in high glucose DMEM (ThermoFisher Scientific, Waltham, MA) plus 10% FetalPlex serum substitute (Gemini Biosciences, West Sacramento, CA) and 1x cell culture Pen/Strep (ThermoFisher Scientific). For transfection, cells were seeded onto 8 well chambered glass coverslips (CellVis, Mountain View, CA; pre-coated with poly-D-lysine) and transfected with eNOS-RFP, Wt HSP90, and/or D88N HSP90, kindly provided by W. Sessa (plasmids # 22497 [RRID:Addgene_22497], #22487 [RRID:Addgene_22487], and/or #22480 [RRID:Addgene_22480], respectively; Addgene, Watertown, MA). S119A eNOS-RFP was generated by site directed mutagenesis from Wt eNOS-RFP and verified by sequencing (University of Pennsylvania Penn Genomic Analysis Core DNA Sequencing Facility). Cells were stimulated with 10 µg/ml SC79 (Cayman Chemical, Ann Arbor MI) that was made from 10mg/ml SC79 stock in DMSO. Control solutions had 0.1% DMSO as vehicle control. No effects of DMSO alone were observed. RFP fluorescence was used to verify similar transfection efficiencies in all experiments.

H441 small airway epithelial cells (American Type Culture Collection [ATCC], Manassas VA; Cat# HTB-174, RRID:CVCL_1561) were cultured in Minimal Essential Media (MEM; ThermoFisher Scientific) with Earle’s salts plus 10% FetalPlex serum substitute and 1x cell culture Pen/Strep. Cells were seeded and imaged on plastic 48 well plates as they stuck very poorly to glass. For air-liquid interface (ALI) cultures, H441 cells were seeded onto collagen-coated 1.1 cm^2^ transwell filters (for intracellular DAF-FM) or 0.33 cm2 transwell filters (for extracellular Daf-2) both with 0.4 µm pore size (transparent; Corning, Corning NY) and grown to confluence for 5 days before exposing the apical side to air, as described previously for 16HBE and Calu-3 cells (20, 67). H441 ALIs were switched to bronchial epithelial cell basal media (BEBM; Lonza, Walkersville MD) plus Lonza Singlequot supplements (differentiation media) upon confluence and exposure to air. When culture medium was removed from the upper compartment, basolateral media was changed to differentiation medium (1:1 DMEM:BEBM) containing hEGF (0.5 ng/ ml), epinephrine (5 ng/ml), BPE (0.13 mg/ml), hydrocortisone (0.5 ng/ml), insulin (5 ng/ml), triiodothyronine (6.5 ng/ml), and transferrin (0.5 ng/ml), supplemented with 100 U/ml penicillin, 100 g/ml streptomycin, 0.1 nM retinoic acid, and 2% NuSerum (BD Biosciences, San Jose, CA) all from the the Lonza Singlequot supplements as described (18). Cells were fed from the basolateral side only with differentiation media for ∼21 days before use.

Submerged H441 cells were infected with Green GENIe-expressing BacMam (Montana Molecular, Bozeman MT) as per the manufacture’s protocol for adherent cells; media was changed 4-6 hours after infection to BacMam-free media containing 2 mM NaButyrate to maintain expression. For siRNA, H441s were treated with Acell SMARTPool siRNAs (Horizon Discovery, Waterbeach, UK) for human eNOS (NOS3; Catalog ID:E-006490-00-0005), nNOS (NOS1; Catalog ID:E-009496-00-0005), PAR-2 (Catalog ID:E-005095-00-0005), or non-targeting pool (Catalog ID:D-001910-10-05) as per the manufacturer’s instructions and as previously described (42). H441 cells were transfected with Wt or D88N HSP90 (Kindly provided by W. Sessa, Addgene plasmids #22487 or #22480, respectively) using Lipofectamine 3000 and the specific H441 protocol provided on the ThermoFisher Scientific website. We used 5 µL of lipofectamine and 2 µg DNA per 8-well chambered coverglass (CellVis), equating to 0.25 µg DNA/well, with each well approximately equivalent to a 48 well plate in surface area.

A549 cells (NCI-DTP Cat# A549, RRID:CVCL_0023) were identically transfected with GFP-eNOS (provided by W. Sessa, Addgene plasmid #22444^;^ RRID:Addgene_22444) and/or mCherry-HSP90 (provided by D. Picard, Addgene plasmid #108223; RRID:Addgene_108223). Cells were cultured in Ham’s F12K media (ThermoFisher Scientific) containing 10% FetalPlex and 1% penicillin/streptomycin mix and imaged in 20 mM HEPES-buffered Hank’s Balanced Salt Solution (HBSS). Cells were transfected and imaged on 8-well chambered coverglass (uncoated) using 470/20 nM band pass (bp) filter (for GFP excitation), 490 lp dichroic beamsplitter, and 520/40 nm bp filter (for GFP emission) or 600/50 nm bp filter (for mCherry emission). Lack of bleed-through of GFP emission with the mCherry emission filter was observed in pilot experiments imaging GFP transfection only and is demonstrated with no fluorescence when GFP-eNOS is expressed with mCherry alone (described in the main text). Imaging of A549 cells was carried out using a 40x 0.75 NA objective on in Olympus (Tokyo, Japan) IX-83 microscope with Hammamatsu (Tokyo, Japan) Orca Flash 4.0 sCMOS camera and XCite 120 Boost LED illumination source (Excelitas, Waltham MA) and MetaMorph (Molecular Devices, San Jose CA).

Primary human M0 macrophages were cultured as previously described (68) in high glucose RPMI2650 media with 10% human serum and 1x cell culture Pen/Strep. De-identified monocytes from healthy apheresis donors were obtained from the University of Pennsylvania Human Immunology core with written informed consent of every participant and institutional review board approval. Cells isolated from 10 different individuals were used. As all samples were de-identified for race, age, sex, etc., samples were used in a blinded fashion. Macrophages were differentiated by adherence culture for 12 days in 8-well chamber slides (CellVis) as described (68). Our prior studies suggest no differences in T2R responses among macrophages differentiated by adherence alone or by adherence plus M-CSF (42), and thus adherence only was used for these studies.

Primary human nasal epithelial cells were obtained in accordance with The University of Pennsylvania guidelines regarding use of residual clinical material from patients undergoing sinonasal surgery at the University of Pennsylvania with institutional review board approval (#800614) and written informed consent from each patient in accordance with the U.S. Department of Health and Human Services code of federal regulation Title 45 CFR 46.116. Inclusion criteria were patients ≥18 years of age undergoing sinonasal surgery for sinonasal disease (CRS) or other procedures (e.g. trans-nasal approaches to the skull base). Exclusion criteria included history of systemic inheritable disease (eg, granulomatosis with polyangiitis, cystic fibrosis, systemic immunodeficiences) or use of antibiotics, oral corticosteroids, or anti-biologics (e.g. Xolair) within one month of surgery. Individuals ≤18 years of age, pregnant women, and cognitively impaired persons were not included. Tissue was transported to the lab in saline on ice and mucosal tissue was immediately removed for cell isolation.

Sinonasal epithelial cells were enzymatically dissociated and grown to confluence in proliferation medium (50% DMEM/Ham’s F-12 plus 50% BEBM plus Lonza Singlequot supplements) for 7 days (18–20, 69). Confluent cells were dissociated and seeded on Corning transwells (0.33 cm2, 0.4 µm pore size; transparent; corning) coated with BSA, type I bovine collagen, and fibronectin (Corning). When culture medium was removed from the upper compartment, basolateral media was changed to differentiation medium as described above for H441 ALIs. Primary ALI cultures were genotyped for *TAS2R38* PAV (functional) or AVI (non-functional) polymorphims (70, 71) as described (18–20). Cell identity was verified based airway epithelial morphology (formation of motile cilia, goblet cells, transepithelial electrical resistance, etc.) observed after differentiation.

### Live cell imaging of Ca^2+^, NO, and cGMP

Unless indicated, all regents were from MilliporeSigma (St. Louis MO). Adherent, submerged HEK293Ts in 20 mM HEPES-buffered Hank’s Balanced Salt solution (HBSS) were simultaneously loaded and stimulated for 30 min in the presence of 10 µM DAF-FM-diacetate (ThermoFisher Scientific) ± SC79 (Cayman Chemical) as indicated. HEK293Ts were then immediately washed three times in HBSS and imaged as below. Submerged H441 cells were loaded for 30 minutes with 10 µM DAF-FM-diacetate (ThermoFisher Scientific) in 20 mM HEPES-buffered HBSS supplemented with 1x MEM amino acids followed by washing with HBSS + 1x MEM amino acids. Calbryte 590 AM (AAT Bioquest, Sunnyvale, CA) was loaded identically. When used, L-NAME (10 µM), D-NAME (10 µM), U73122 (1 µM), U73343 (1 µM), or geldanamycin (all from Cayman Chemical) were included in the loading solution as 30 min pretreatment; cPTIO (10 µM) was only added after loading. Cells were then washed out of

DAF-FM into HBSS with the continued presence of inhibitor for the start of the experiment. Transwells were loaded with DAF-FM-diacetate for 60 min and placed into a glass-bottom 12 well dish (CellVis) prior to imaging. Macrophages were loaded with 5 µM fura-2-AM or DAF-FM DA for 45 min as previously described (68). Primary human ALIs were loaded for 90 min with DAF-FM-diacetate as previously described (18, 19, 69). Denatonium benzoate, sodium benzoate, and phenylthiocarbamide were from Sigma Aldrich (St. Louis, MO USA) and N-(acetyloxy)-3-nitrosothiovaline SNAP, BIIB021, and VER-155008 were from Cayman Chemical (St. Louis, MO USA) DAF-FM, fura-2, and cGMP were imaged as previously described (20, 68). DAF-FM was imaged on a TS100 microscope (20x 0.75 PlanApo objective for macrophages on glass and 10x 0.3 NA PlanFluor objective for H441 cells submerged on plastic or grown on transwells; Nikon, Tokyo, Japan) GFP filter set, XCite 110 LED (Excelitas Technologies, Waltham MA USA), and Retiga R1 Camera (Teledyne QImaging, Surrey, BC, Canada). Calbryte 590 was imaged using the same microscope plus TRITC filter set and 10x 0.3 NA PlanFluor objective, as submerged H441s on plastic necessitated a longer working distance. Images were acquired using Micromanager (72). Fura-2 was imaged using MetaFluor (Molecular Devices, Sunnyvale, CA USA) and standard fura-2 dual excitation filter set on IX-83 microscope (20x 0.75 NA PlanApo objective for macrophages on glass, 10x 0.4 NA PlanApo objective for H441 cells on plastic; Olympus, Tokyo, Japan) equipped with a fluorescence xenon lamp (Sutter Lambda LS, Sutter Instruments, Novato, CA USA), excitation and emission filter wheels (Sutter Lambda 2), and Orca Flash 4.0 sCMOS camera (Hamamatsu, Tokyo, Japan). Green GENIe cGMP construct was imaged using a FITC filter set, IX83 microscope, 10x 0.4 NA PlanApo objective, XCite 120Boost LED illumination, and MetaMorph.

### Measurement of ciliary beat frequency (CBF)

Whole-field CBF was measured using the Sisson-Ammons Video Analysis system (73) as previously described (18, 74, 75) at ∼26-28 °C, with the exception of bacterial cHBSS experiments, which were carried out at room temperature. Cultures were imaged at 120 frames/second using a Leica DM-IL microscope (20x/0.8 NA objective) with Hoffman modulation contrast in a custom glass bottom chamber. Experiments utilized Dulbecco’s PBS (+ 1.8 mM Ca^2+^) on the apical side and 20 mM HEPES-buffered Hank’s Balanced Salt Solution supplemented with 1× MEM vitamins and amino acids on the basolateral side. As typically done with CBF measurements (74–77), changes in CBF were normalized to baseline CBF. This was validated by measurements of raw baseline CBF (in Hz) between control and experimental cultures showing no significant differences, as indicated in the text.

### Bacteria culture

PAO-1 (ATCC 15692) and PAO-JP2 (ΔlasI, ΔrhlI; Tc^r^, HgCl_2_^r^; (78, 79) were cultured in LB media as described (18, 67). Conditioned HBSS (cHBSS) was prepared by taking the pellet of an overnight culture and resuspending to OD 0.1 in HBSS and incubating overnight with shaking. We used cHBSS over conditioned LB due to the slight stimulatory effects of LB alone on CBF at dillutions >10% (18). After centrifuging (5000 x g, 15 min, 4°C) to pellet bacteria, cHBSS was filtered through a 0.2 µm filter then diluted as indicated with unconditioned (unmodified) HBSS.

### Immunofluorescence (IF) microscopy

IF was carried out as previously described (18–20). ALI cultures were fixed in 4% formaldehyde for 20 min at room temperature, followed by blocking and permeabilization in phosphate buffered saline (PBS) containing 1% bovine serum albumin (BSA), 5% normal donkey serum (NDS), 0.2% saponin, and 0.3% triton X-100 for 1 hour at 4°C. A549 or 16HBE cells were fixed in 4% formaldehyde for 20 min at room temp, followed by blocking and permeabilization in PBS containing 1% BSA, 5% NDS, 0.2% saponin, and 0.1% triton X-100 for 30 min at 4°C. Primary antibody incubation (1:100 for anti-T2R antibodies, 1:250 for tubulin antibodies) were carried out at 4°C overnight. Incubation with AlexaFluor (AF)-labeled donkey anti-mouse and rabbit secondary antibody incubation (1:1000) was carried out for 2 hours at 4°C. Transwell filters were removed from the plastic mounting ring and mounted with Fluoroshield with DAPI (Abcam; Cambridge, MA USA)). For co-staining of T2R14 and T2R38, Zenon antibody labeling kits (Thermo Fisher Scientific) were used to directly label primary antibodies with either AF546 or AF647 as described (18–20). Images of ALIs were taken on an Olympus Fluoview confocal system with IX-73 microscope and 60x (1.4 NA) objective and analyzed in FIJI (80). Images of submerged H441 cells were taken on an Olympus IX-83 microscope with 60x (1.4 NA) objective using Metamorph. Anti-T2R38 (ab130503; rabbit polyclonal; RRID:AB_11156286) and anti-beta-tubulin IV (ab11315; mouse monoclonal; RRID:AB_297919) antibodies were from Abcam. Anti-T2R14 (PA5-39710; rabbit polyclonal; RRID:AB_2556261) primary antibody and conjugated secondary antibodies (donkey anti-rabbit AlexaFluor 546 [RRID:AB_2534016] and donkey anti-mouse AlexaFluor 488 [RRID:AB_141607]) were from ThermoFisher Scientific. Alpha-tubulin antibody was from Developmental Studies Hybridoma Bank (12G10; mouse monoclonal; University of Iowa, Iowa City; RRID:AB_1157911). Anti-eNOS antibody (NB-300-605; rabbit polyclonal; RRID:AB_10002794) was from Novus (Littleton, CO). Immunofluorescence images were analyzed in FIJI (80) using only linear adjustments (min and max), set equally between images that are compared. Compared images were always taken with the same exposure, objective, and other camera and microscope settings. Both conventional (0 = black) and inverted (0 = white) lookup tables (LUTs) were used in this study to illustrate localizations as clearly as possible. Inverted LUTs were from ChrisLUTs FIJI package (https://github.com/cleterrier/ChrisLUTs; C. Leterrier, Neuropathophysiology Institute, Marseille University).

### Phagocytosis assays

Phagocytosis assay were carried out as descried (68). Macrophages were incubated with heat-killed FITC-labeled *Escherichia coli* at 250 µg/ml (strain K-12; reagents from Vybrant phagocytosis assay kit; ThermoFisher Scientific; Cat # E2861) in phenol red-free, low glucose DMEM (ThermoFisher Scientific) ± denatonium benzoate or other agonists or inhibitors for 15 min at 37°C. As we found that phagocytosis was negligible from temperatures of 4 °C up to room temp in these assays (42), we recorded fluorescence from living cells at room temperature immediately after the 15 min 37 °C incubation with FITC-*E. coli*. Extracellular FITC was quenched with trypan blue per the manufacturer’s instructions, and fluorescence was recorded on a Spark 10M plate reader (Tecan; 485 nm excitation, 535 nm emission). For representative micrograph shown, macrophages on glass were incubated as above, and extracellular FITC was quenched with trypan blue and cells were washed ≥5x in PBS to remove residual extracellular FITC-*E. coli*. Remaining adherent MΦs were fixed in 4% formaldehyde (Electron Microscopy Sciences, Hatfield, PA) for 10 min followed by DAPI staining in mounting media (Fluoroshield with DAPI, Abcam). FITC-E. coli were then imaged using standard FITC filter set (Semrock, Rochester, NY) on an inverted Olympus IX-83 microscope with 20x (0.8 NA) objective, XCite 120LEDBoost illumination source, and Hammamatsu Orca Flash 4.0 sCMOS camera.

Phagocytosis assays were also carried out similarly using 125 µg/ml pHrodo red-labeled *S. aureus* (strain Wood 46; ThermoFisher Scientific, cat # A10010) (68). As pHrodo dyes only fluoresce when particles are internalized into low pH endosomes (previously demonstrated in (42)), this assay does not require washing or quenching of the extracellular pHrodo *S. aureus*. Macrophages were incubated with pHrodo-*S. aureus* for 30 min at 37°C as described (68) with excitation at 555 nm and emission at 595 nm measured on the Tecan Spark 10M plate reader. Background measurements were made using wells containing fluorescent *S. aureus* in the absence of macrophages. Representative images were taken as above except using a standard TRITC filter set (Semrock).

### Data analysis and statistics

Multiple comparisons were made with one-way ANOVA with Bonferroni (pre-selected pairwise comparisons), Tukey-Kramer (comparing all values), or Dunnett’s (comparing to control value) post-tests; *p* <0.05 was considered statistically significant. Asterisks (* and **) indicate *p* <0.05 and *p* <0.01, respectively. All data in bar graphs are shown as mean ± SEM with n derived from biological replicates (separate experiments conducted with different passage/patient cells on different days). Images shown for comparison were collected on the same day under identical conditions with identical min/max settings. No non-linear (e.g., gamma) adjustments were made to any images for either display or analysis. Raw unprocessed image data were analyzed in FIJI (80) and resulting numerical data were analyzed in Excel (Microsoft) and/or Prism (GraphPad software, La Jolla, CA). All data used to generate bar graphs and traces are available upon request.

## RESULTS

### HSP90 inhibition reduces heterologously-expressed eNOS function in HEK293Ts and A549s

To first determine if we could recapitulate results that HSP90 is important for eNOS function (62, 81–84) in a reductionist model, we expressed eNOS-RFP in HEK293Ts, an eNOS null cell line (85). We measured NO production using reactive nitrogen species (RNS)-sensitive dye DAF-FM over 30 min. One mechanism by which eNOS can be activated is phosphorylation at S1177 (S1179 in bovine eNOS). We found that expression of S1179D eNOS dramatically increased DAF-FM fluorescence compared with Wt eNOS-RFP or S1179A eNOS-RFP (**Figure 1A-B**). DAF-FM fluorescence increases likely reflected NO production as they were inhibited after 30 min pre-treatment and in the continued presence of 10 µM L-NAME but not inactive control D-NAME (**Figure 1B**). We also observed that HSP90 inhibitor geldanamycin (10 µM; 30 min pre-treatment then continued throughout the 30 min experiment) also reduced DAF-FM fluorescence in S1179D eNOS-RFP-expressing cells. Supporting that the effects of geldanamycin were mediated by HSP90 inhibition, we also found that co-transfection of a dominant negative (DN) HSP90 isoform (D88N) (84) reduced DAF-FM fluorescence while Wt HSP90 did not significantly change DAF-FM fluorescence (**Figure 1B**).

**Figure 1.**
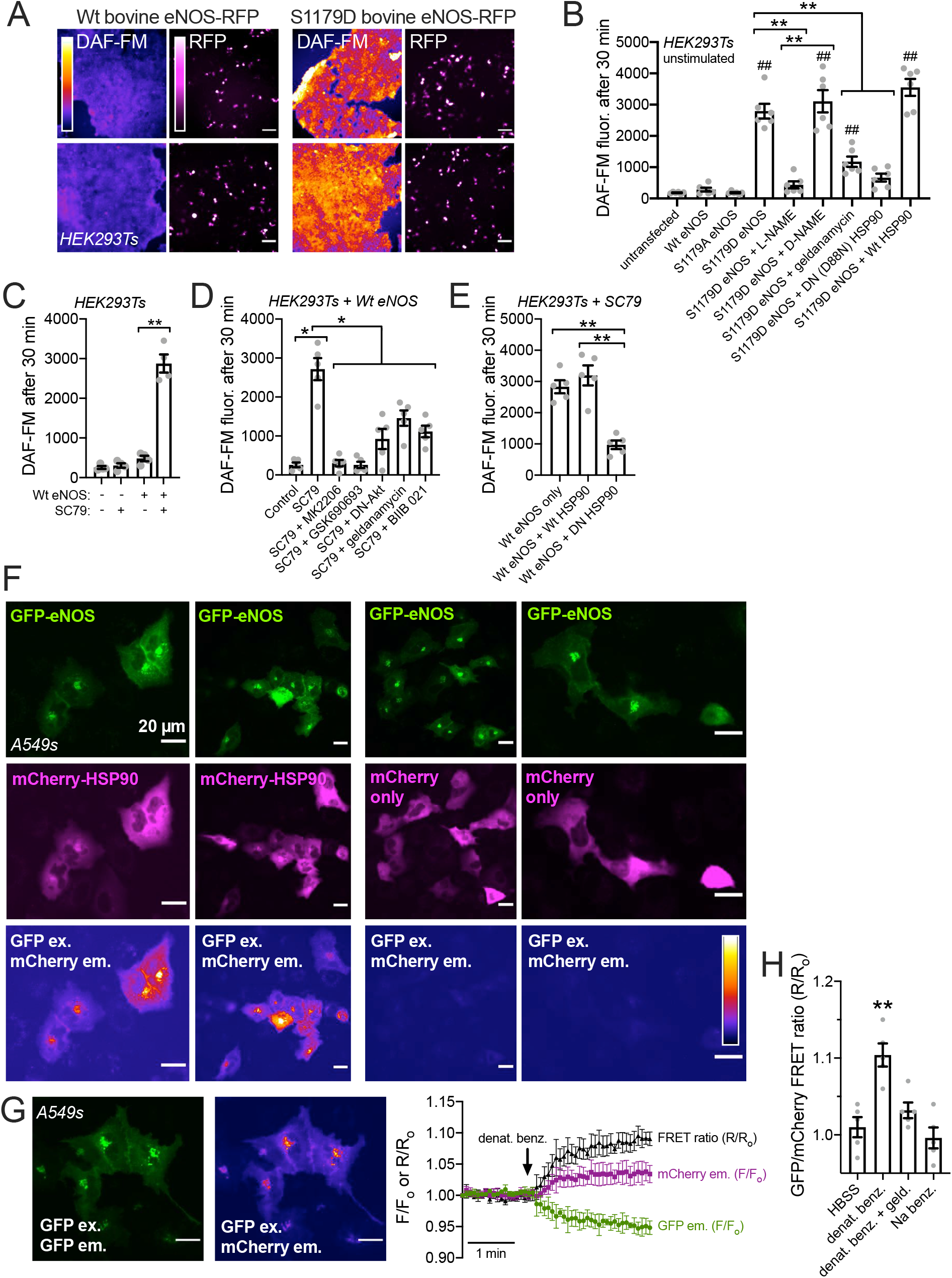
Role of HSP90 in NO production by heterologously-expressed endothelial nitric oxide synthase (eNOS) in HEK293T and A549 cells. ***A*:** Live-cell images of HEK293T cells expressing bovine eNOS-RFP (either wild type [Wt] or S1179D) after 30 min loading with DAF-FM. ***B:*** Bar graph of cell fluorescence intensity from experiments as in B. S1179D eNOS expression increased DAF-FM fluorescence over Wt or S1179A eNOS. DAF-FM fluorescence increase with S1179D eNOS was inhibited by NOS inhibitor L-NAME, HSP90 inhibitor geldanamycin, or dominant negative (DN) D88N HSP90. Bar graph is mean ± SEM of n = 6 independent experiments, shown by data points. Significance by Bonferroni posttest; \*\**p*<0.01 vs bracketed groups and ^##^*p*<0.01 vs transfected control. **C:** In HEK293Ts expressing Wt eNOS-RFP, SC79 increased DAF-FM fluorescence after 30 min loading. Bar graph shows mean ± SEM of 4 independent experiments. Significance by Bonferroni posttest comparing untransfected ± SC79 and comparing eNOS transfected ± SC79; ** *p*<0.01. ***D:*** In HEK293Ts expressing Wt eNOS, SC79 increased DAF-FM fluorescence after 30 min loading, which was reduced with Akt inhibitors MK2206, GSK690693, co-transfection of dominant negative (DN)-Akt, or HSP90 inhibitors geldanamycin or BIIB 021. Bar graph shows mean ± SEM (n = 5 independent experiments). Significance by Bonferroni posttest with paired comparisons; \**p*<0.05. ***E:*** In HEK293Ts stimulated with SC79 and transfected with Wt eNOS, co-transfection of DN HSP90 but not Wt HSP90 reduced DAF-FM fluorescence after 30 min loading. Bar graph shows mean ± SEM (n = 5 independent experiments). Significance by one way ANOVA with Bonferroni posttest; \*\**p*<0.01. ***F:*** Representative images (3 independent transfection experiments) of A549 cells transfected with GFP-eNOS (green) and mCherry HSP90 (magenta; left 2 columns) or mCherry alone (magenta; right 2 columns). Intensity pseudocolored images at the bottom show fluorescence with GFP excitation filters and mCherry emission filter, shown at identical microscope settings for all 4 columns. ***G:*** Representative image showing 3 A549 cells transfected with GFP-eNOS and mCherry-HSP90 imaged with GFP filters (left) or GFP excitation and mCherry emission filters (right). Trace on the right shows normalized (F/F°) average of GFP vs mCherry emission (both using GFP excitation) as well as the ratio of mCherry/GFP emission (FRET ratio; R/R°). Note that 1 mM denatonium benzoate increases mCherry (acceptor) emission but decreases GFP (donor) emission, suggesting a bona fide increase in FRET. ***H:*** Bar graph showing mean ± SEM of R/R° from independent experiments (n = 5) as in G. Cells were stimulated with HBSS only (control), sodium benzoate (1 mM), or denatonium benzoate (1 mM) ± geldanamycin (10 µM; 30 min pre-treatment plus stimulation with continued 10 µM geldanamycin). HBSS, denat. benz., and Na benzoate conditions contained 0.1% DMSO as vehicle control. Significance by one way ANOVA with Dunnett’s posttest comparing all values to HBSS alone; \*\**p*<0.01.

Small molecule Akt activator SC79 also induces eNOS phosphorylation and NO production in airway epithelial cells (86). We found that SC79 (10 µg/ml) activates DAF-FM fluorescence increases in HEK293Ts transfected with Wt eNOS but not untransfected cells (**Figure 1C**). In Wt eNOS-transfected cells, SC79-induced DAF-FM fluorescence increases were reduced by Akt inhibitor MK2206 or GSK690693 (10 µM; 30 min pre-treatment then continued throughout the 30 min experiment) as well as co-transfection with dominant negative (K179M) Akt (87) (**Figure 1D**). SC79-induced DAF-FM fluorescence increases were also blocked by HSP90 inhibitors geldanamycin or BIIB 021 (**Figure 1D**). Further supporting the role of HSP90 in eNOS function, we found that co-transfection of DN HSP90 in SC79-stimulated HEK293Ts reduced DAF-FM fluorescence (**Figure 1E**). All together, these data suggest that HSP90 is important for eNOS-mediated NO production.

When we transfected GFP-tagged eNOS (88) and mCherry-tagged HSP90 (89) into submerged A549 airway cells, chosen here because of their transfectability and their ability to stick well to glass. We saw punctate perinuclear localization of eNOS likely reflecting Golgi, as eNOS localizes partly to the Golgi in endothelial cells (85) (**Figure 1F**). We also saw some likely plasma membrane eNOS localization at cell-cell contact points (**Figure 1F**), also expected from studies of endothelial cells (88). HSP90 localization was more global (**Figure 1F**), though perinuclear puncta were observed. However, when we exited GFP and collected mCherry emission, we saw an mCherry emission signal that almost identically overlapped with GFP-eNOS localization (**Figure 1F**). As GFP and mCherry are a donor-acceptor pair for Förster resonance energy transfer (FRET), we hypothesized that we were collecting emission from mCherry-HSP90 in close proximity to the excited GFP-eNOS. When this experiment was performed with GFP-eNOS and mCherry alone (no HSP90), no mCherry emission was detected with GFP excitation (**Figure 1F**). A549 cells endogenously express T2R bitter taste receptors activated by the bitter agonist denatonium benzoate (90). When we monitored GFP and mCherry emission, both with GFP excitation, we noted an increase in mCherry emission and concomitant decrease in GFP emission with denatonium benzoate stimulation (**Figure 1G**), suggesting that an increase in FRET occurs with bitter agonist stimulation. This increase in FRET was not observed during stimulation with Na benzoate and was reduced with HSP90 inhibitor geldanamycin (**Figure 1H**). Together, these A549 data suggest that heterologously-expressed HSP90 and eNOS are partly closely co-localized in an airway cell line, and this association or close co-localization may increase during T2R stimulation.

### HSP90 inhibition reduces endogenous eNOS function in H441 cells

We next wanted to test if HSP90 activity affects endogenous eNOS function when activated by endogenous T2R receptors. We started by examining if T2R stimulation activates NO production in H441 small airway epithelial cells, a club cell-like cell line that expresses eNOS similarly to primary bronchial cells (31, 91). H441 cells produce NO in response to estrogen and other types of stimulation (31, 91–94). We observed positive immunofluorescence (IF) for eNOS in submerged H441s compared with rabbit serum and fluorescent secondary alone (**Figure 2A-B**). This eNOS signal was blocked by pre-treatment of H441 cells with eNOS siRNA (**Figure 2C**), confirming that H441s express eNOS as demonstrated previously by others using Western (31, 91).

**Figure 2.**
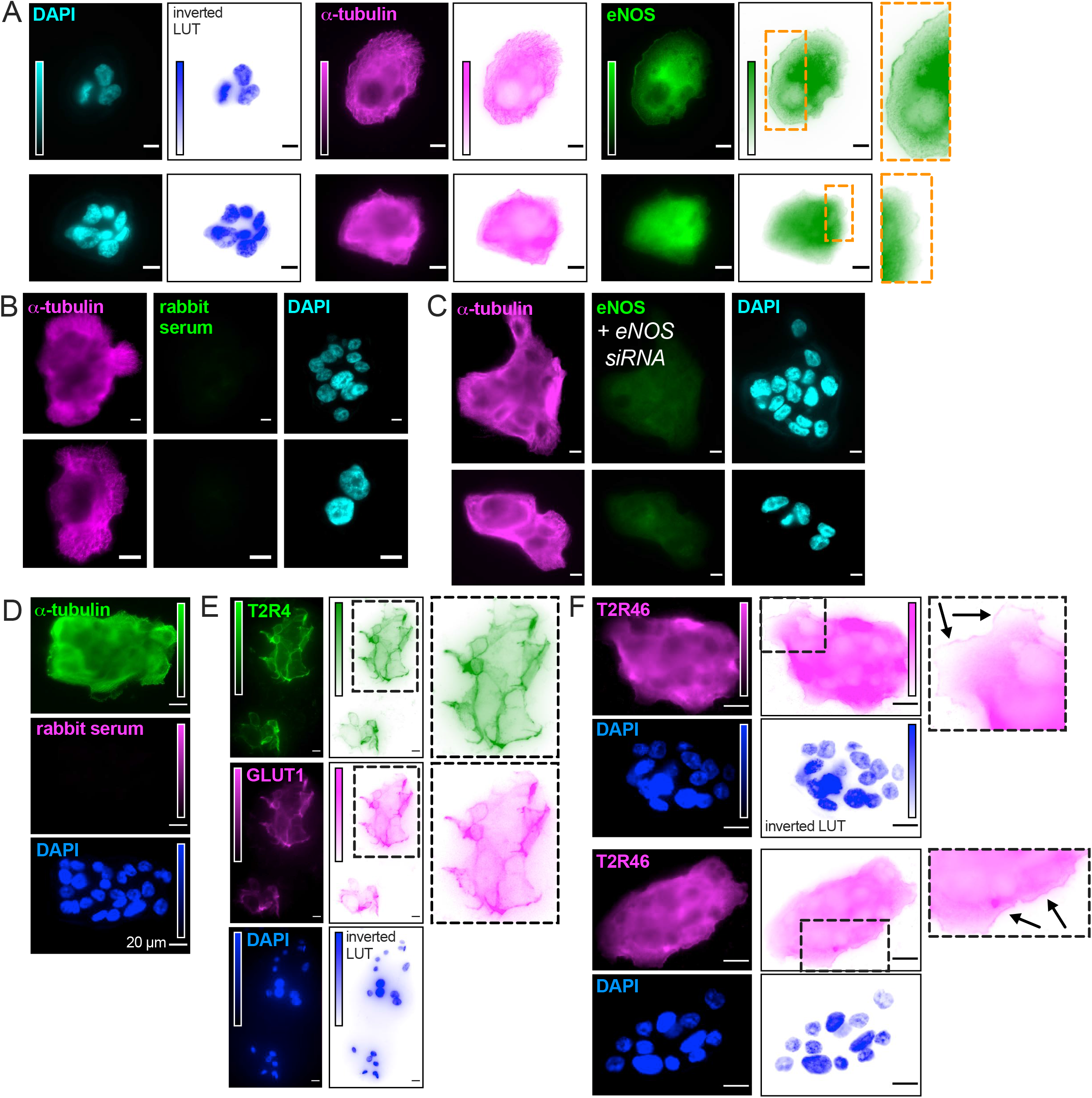
eNOS, T2R4, and T2R46 immunofluorescence in submerged H441 cells. ***A*:** Immunofluorescence images of cells stained with primary mouse monoclonal antibody for α-tubulin and rabbit polyclonal antibody to eNOS. Inverted look-up table (LUT) images are shown along with conventional LUTs to highlight localization. ***B*:** Rabbit serum plus secondary antibody control. Cells were stained, fixed, and incubated in parallel with cells from experiments as in (A), and imaged at identical microscope settings (objective, exposure time, *etc.*). ***C*:** Images of eNOS siRNA-treated cells stained with primary mouse monoclonal antibody for α-tubulin and rabbit polyclonal antibody to eNOS, imaged at identical microscope settings to A and B. ***D-E***: Images of rabbit serum only (D) or T2R4 (E) localization, showing similar pattern of T2R4 and GLUT1 staining (D). ***F*:** T2R46 stained cells with inset regions blown up to show membrane-like staining at the edges of the epithelial cell patches. All images representative of ≥3 independent experiments. Scale bars in A-C are 10 µm. Scale bars in D-F are 20 µM.

We also noted positive T2R4 and T2R46 immunofluorescence in submerged H441s (**Figure 2D-F**). T2R4 staining was highly similar to the pattern observed for plasma membrane glucose transporter Glut1 (**Figure 2E**), while T2R46 appeared to be more diffusely localized but also possibly localized to the edges of H441 cell islands (**Figure 2F**). Like many GPCRs, a substantial amount of T2R46 immunofluorescence was located intracellularly, possibly representing ER and/or trafficking compartments. However, potential plasma membrane staining was observed at the cell periphery. The implications for these different staining patterns are unclear, but our goal here was to test for T2R expression and not perform detailed localization analysis. The rationale for examining T2R4 and 46 was that T2R4 localizes to nasal cilia (19, 20) and T2R46 localizes to bronchial cilia (5). Both are also expressed in human monocyte-derived macrophages (42). Immunofluorescence appears to support that both T2R4 and T2R46 are also expressed in H441s in addition to downstream eNOS. Quantitative PCR (qPCR) of submerged H441s for the T2Rs responsive to denatonium benzoate also suggested expression of both T2R4 and T2R46, as well as T2R30 (formerly known as T2R47) and possibly T2R13 and T2R10 (**Figure 3A**).

**Figure 3.**
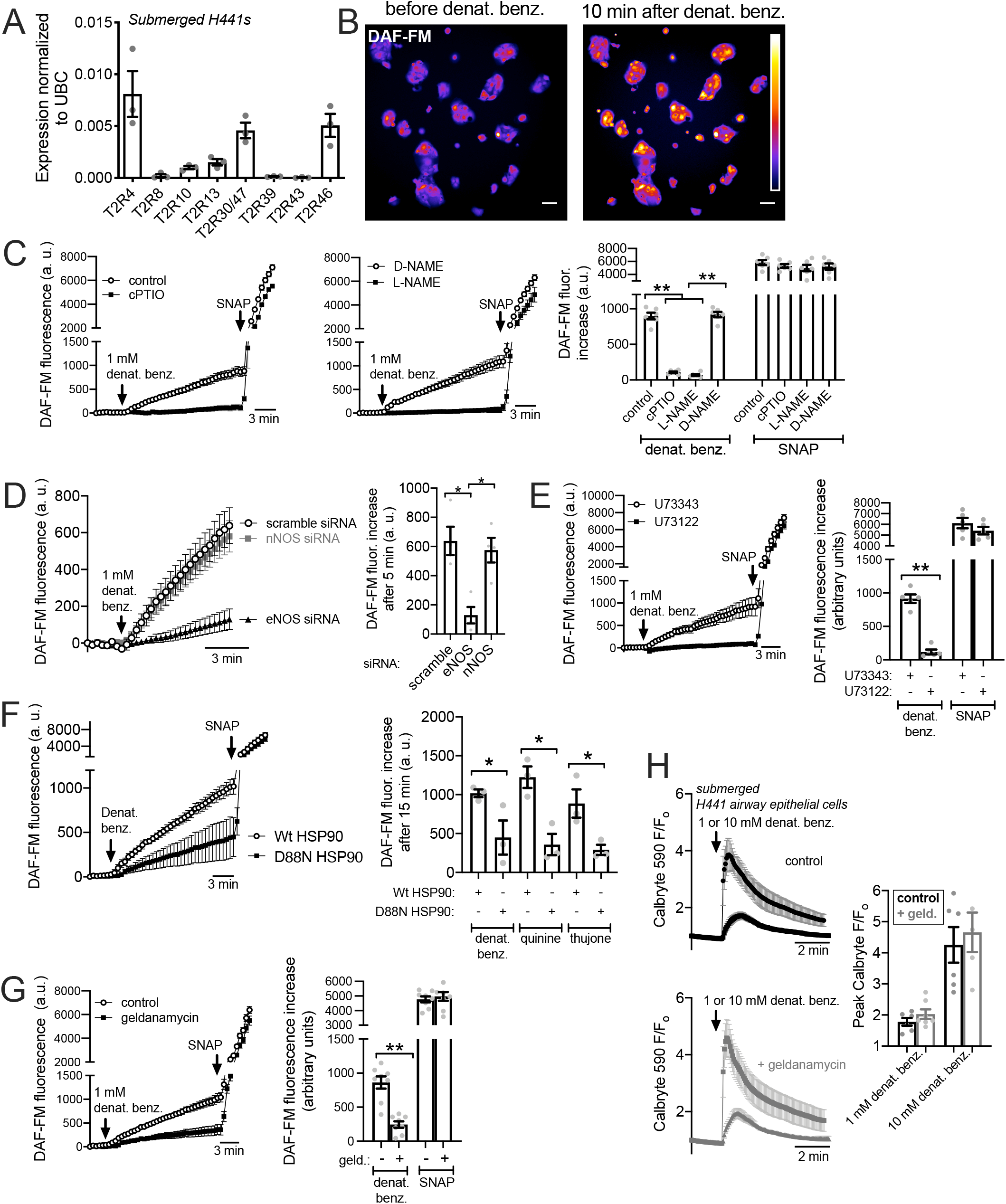
T2R agonist denatonium benzoate activates HSP90-dependent NO production in submerged H441. ***A:*** qPCR of denatonium-responsive *TAS2R* gene expression in H441 cells compared to UBC expression. Bar graph shows mean ± SEM of n = 3 experiments. ***B:*** Representative images of DAF-FM-loaded subconfluent H441 cells immediately before and 10 min after stimulation with 1 mM denatonium benzoate. Each distinct cell island was treated as one region of interest within one experiment. Scale bar is 20 µm. ***C:*** Left and middle show representative traces from H441 cells stimulated with 1 mM denatonium benzoate (denat. benz.) in the presence or absence of NO scavenger cPTIO, NOS-inhibitor L-NAME, or inactive D-NAME. Right is bar graph of mean ± SEM of DAF-FM fluorescence increase after 10 min from 5-8 independent experiments. Significance by Bonferroni posttest with paired comparisons; \*\**p*<0.01. ***D*:** Experiments as in C but done in cells treated with eNOS (NOS3) or nNOS (NOS1) siRNA. Bar graph is mean ± SEM of DAF-FM fluorescence increase after 5 min from 5-8 independent experiments. Significance by Bonferroni posttest; \**p*<0.05. ***E*:** Representative DAF-FM experiments (left; each trace from single experiment showing mean ± SEM of 6-8 H441 cells regions stimulated with 1 mM denatonium benzoate [denat. benz.]) and bar graph (right; mean ± SEM) from experiments in the presence of PLC inhibitor U73122 or inactive control U73343. Bar graph shows results of DAF-FM fluorescence increases after 10 min from 5-8 independent experiments per condition. Significance by Bonferroni posttest with paired comparisons; \*\**p*<0.01. ***F***: Representative DAF-FM traces (left and bar graph) showing stimulation of H441 cells after transfection with Wt or dominant negative (D88N) HSP90. Significance in bar graph by Bonferroni posttest with paired comparisons; \**p*<0.05. ***G*:** Representative experiment (left) and bar graph (right) from experiments as in *B-E* but done ± HSP90 inhibitor geldanamycin. Significance by Bonferroni posttest with paired comparisons; \*\**p*<0.01. ***H:*** Left and middle shows representative Ca^2+^ responses during denatonium stimulation in the absence (left) or presence (right) of geldanamycin, imaged using Calbryte 590. Bar graph shows responses from 5-7 independent experiments per condition. No significant difference determined by determined by one-way ANOVA plus Bonferroni posttest with paired comparisons.

All of these T2Rs (4, 46, 10, 13, and 30) are activated by the bitter compound denatonium benzoate, which activates eight out of 25 human T2R isoforms (95, 96). The denatonium benzoate effective concentrations for T2R4 and T2R46 are 300 and 30 µM, respectively, with an EC50 of ∼240 µM reported for T2R46 in heterologous HEK293T expression systems. To test if denatonium benzoate activated NO production in submerged H441s, we loaded these cells with reactive nitrogen species-sensitive dye DAF-FM to track NO production, as done previously in primary nasal cells (19, 20). Stimulation of DAF-FM-loaded H441s with 1 mM denatonium benzoate caused an increase in intracellular DAF-FM fluorescence (**Figure 3B**). The S-nitrosothiol NO donor SNAP (20 µM) was used in these experiments as a positive control. Denatonium-induced DAF-FM responses were also inhibited by NO scavenger carboxy-PTIO (cPTIO; 10 µM added at the beginning of the experiment; **Figure 3C**) or after 45 min pre-treatment with 10 µM NOS inhibitor L-NAME but not inactive D-NAME (**Figure 3C**). Furthermore, H441 DAF-FM responses to denatonium benzoate were inhibited by eNOS but not nNOS or scramble siRNA (**Figure 3D**). This supports that the DAF-FM responses measured reflect eNOS-mediated NO production.

The denatonium-induced DAF-FM fluorescence increases were inhibited by phospholipase C (PLC) inhibitor U73122 (10 µM; 45 min pre-treatment) but not inactive analogue U73343 (**Figure 3E**), as observed in primary nasal cells (19, 20) and macrophages (42). This suggests the DAF-FM responses require PLC IP^3^ generation and Ca^2+^ signaling, likely downstream of T2R GPCR activation.

We also observed that HSP90-induced DAF-FM increases were reduced in cells transfected with a dominant negative HSP90 beta (D88N-HSP90; **Figure 3F**). D88N-HSP90 blocks VEGF-induced NO production in endothelial cells (84). Transfection of Wt HSP90 had no effect on the denatonium response (**Figure 3F**). We also saw reduction of NO responses with D88N-HSP90 during stimulation with T2R agonists quinine (500 µM) and thujone (1 mM) (**Figure 3F**), which activate T2Rs in bronchial (5) and nasal cilia (97). Quinine activates 11 T2Rs including T2R4 and 46, while Thujone activates T2Rs 14 and 10 (96, 98). These results suggest that T2R agonist-stimulated NO responses require HSP90 function in H441 cells. We also saw that denatonium-induced DAF-FM fluorescence increases were also reduced ≥50% by pretreatment (1 hour) with 10 µM HSP-90 inhibitor geldanamycin (**Figure 3G**), suggesting denatonium-induced (likely T2R-induced) NO production requires HSP90 activity. There was no alteration of the denatonium-induced Ca^2+^ response with this concentration of geldanamycin (**Figure 3H**), suggesting the role of HSP90 is likely downstream of the Ca^2+^ signaling.

NO increases intracellular cGMP by activating soluble guanylyl cyclase. We tested if denatonium stimulation increased cGMP using a cGMP fluorescent biosensor (Green GENie; Montana Molecular). Denatonium caused an increase in cGMP (decrease in Green GENIe F/F°; plotted inversely in **Figure 4A-B**) that was likewise inhibited by blockade of NOS activity by L-NAME (**Figure 4A and C**) or geldanamycin pretreatment (**Figure 4B and C**). These results all suggest that full activation of eNOS and downstream cGMP production during T2R agonist stimulation in H441 cells requires HSP90 function.

**Figure 4.**
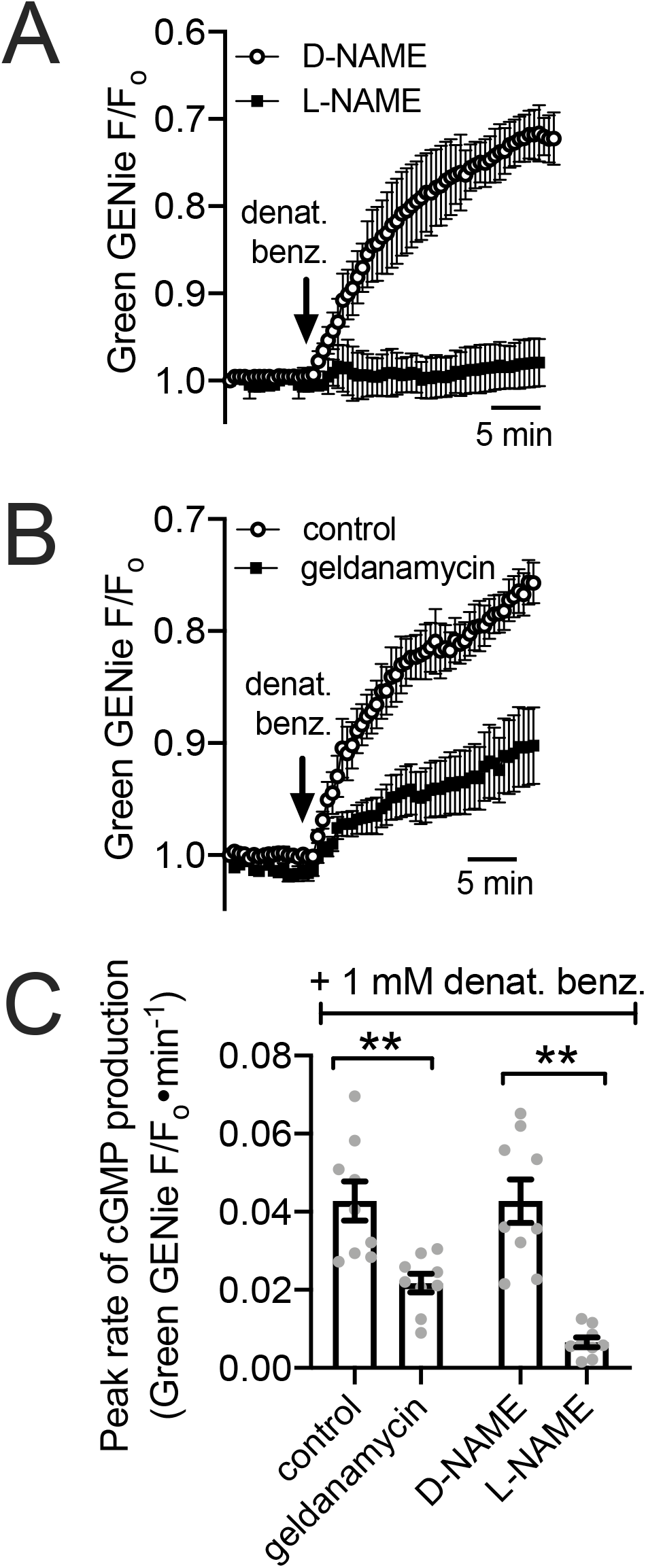
Geldanamycin inhibits NOS-dependent cGMP responses during denatonium benzoate stimulation in H441 cells. ***A:*** Left and middle show traces of Green GENIe cGMP biosensor experiments in H441 cells during denatonium stimulation ± D-NAME or L-NAME. Traces are plotted inversely as F/F° goes down when cGMP goes up; thus, an upward deflection on the trace is an increase in cGMP. ***B:*** Left and middle show traces of Green GENIe cGMP biosensor experiments in H441 cells during denatonium stimulation ± geldanamycin. Control cultures were pre-treated with 0.1% DMSO as a vehicle control. ***C:*** Bar graph quantifying mean ± SEM from 6-8 independent experiments. Significance by Bonferroni posttest with paired comparisons; \*\**p*<0.01.

We grew H441 cell monolayers at ALI, a more physiological cell culture model for airway epithelial cells than submerged cells. H441 ALIs have been used to study ion transport and other epithelial parameters (99–103). Denatonium benzoate, but not sodium benzoate, activated NO production in H441 ALIs, measured by intracellular DAF-FM fluorescence (**Figure 5A-B**). While sodium benzoate does activate some T2Rs, it activates a subset (T2R14 and T2R16) distinct from denatonium benzoate with much lower affinity (96). Thus, we use sodium benzoate as a control here for osmotic effects and potential pH effects due to permeation of the benzoate moiety. Denatonium-induced DAF-FM fluorescence increases were completely blocked by Ca^2+^ chelation by intracellular BAPTA-loading (45 min, 10 µM) plus 0-Ca^2+^ extracellular buffer (containing 2 mM EGTA to chelate trace calcium; **Figure 5C**). This suggests a requirement for Ca^2+^ signaling. Like submerged cells, the DAF-FM response was also reduced by 45 min pre-treatment with HSP90 inhibitor geldanamycin (10 µM) or NOS inhibitor L-NAME (10 µM) but not HSP70 inhibitor VER-155008 (10 µM; **Figure 5D**). Thus, HSP90 inhibition reduces T2R-mediated NO production in H441 ALIs.

**Figure 5.**
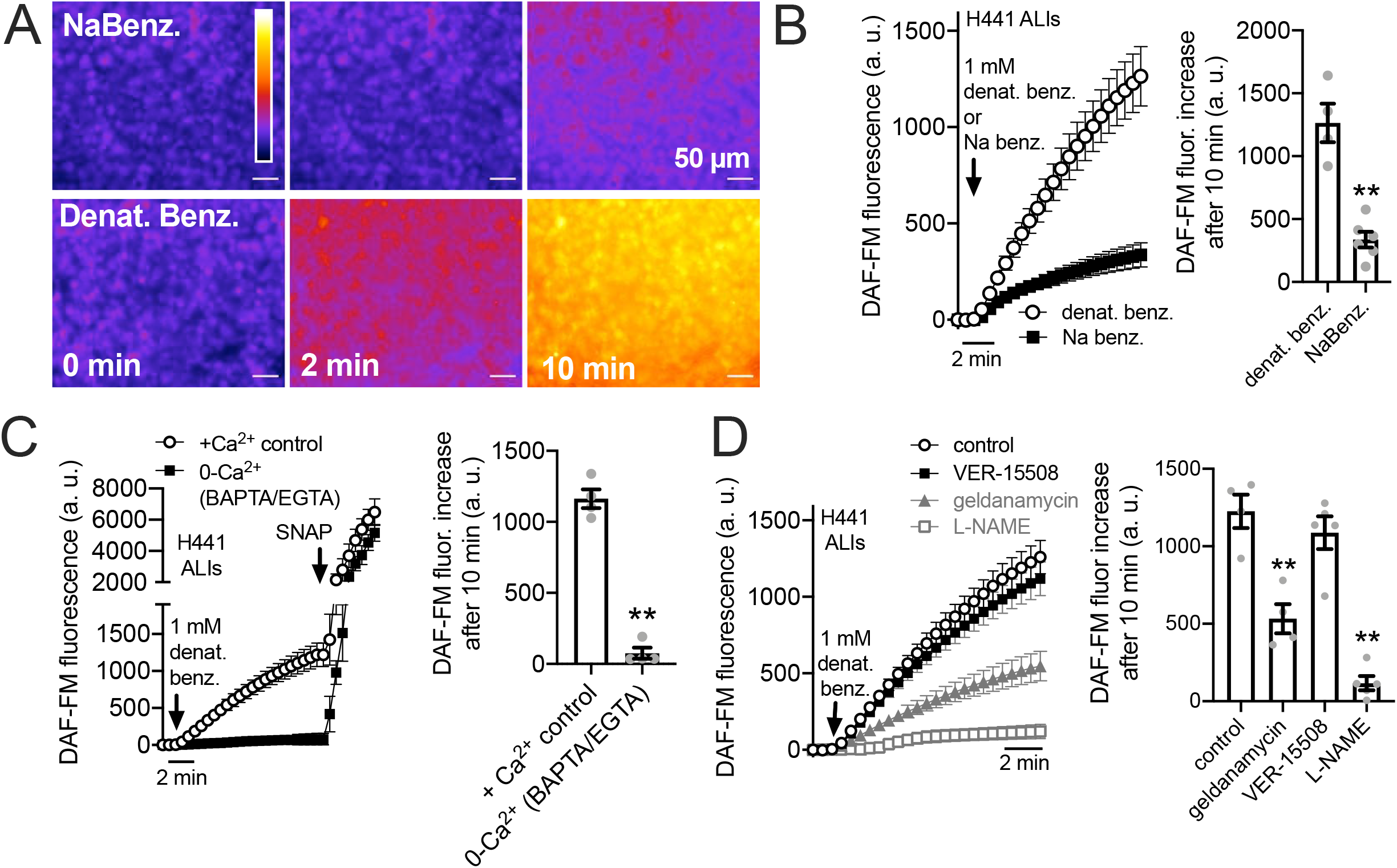
HSP90 inhibition reduces T2R-stimulated intracellular NO production in H441 cells grown at air liquid interface (ALI). ***A:*** Representative image of DAF-FM-loaded H441 ALIs stimulated for 10 min with 1 mM sodium benzoate (NaBenz.) or denatonium benzoate (denat benz.); fluorescence increased with denatonium benzoate but not sodium benzoate. ***B:*** Average trace and bar graph (mean ± SEM) of 4 experiments as in a. Significance determined by Student’s t test; \*\**p*<0.01. ***C:*** Average trace and bar graph (mean ± SEM of 3 experiments) showing response in cultures pre-loaded with BAPTA-AM and stimulated in the absence of extracellular Ca^2+^ (0-Ca^2+o^) vs control cultures pre-incubated with 0.1% DMSO only and stimulated in the presence of extracellular Ca^2+^. ***D:*** Denatonium-induced DAF-FM fluorescence increases in H441 ALIs increases was inhibited by pretreatment with geldanamycin or L-NAME but not HSP70 inhibitor VER-15508. Shown is average trace and bar graph of results from 4 independent experiments. Significance determined by 1-way ANOVA with Dunnett’s posttest comparing all values to control (denatonium only); \*\**p*<0.01.

NO is a highly diffusive gas that can rapidly diffuse across cell membranes (104). When a small volume (100 uL) of cell impermeant NO indicator DAF-2 (10 µM) was placed on top of the H441 ALIs in the presence of denatonium benzoate or sodium benzoate for 30 min, we observed a 3-fold higher fluorescence of apical DAF-2 in denatonium-treated cultures (**Figure 6A**). This suggest that NO produced can diffuse into the airway surface liquid, as previously show in primary sinonasal ALIs (18). The denatonium-induced DAF-2 fluorescence increase was reduced by PLC inhibitor U73122 but not inactive U73343 (**Figure 6A**), suggesting it depended on GPCR signaling. We found that T2R agonists denatonium benzoate and quinine, but not sodium benzoate, both increased apical DAF-2 fluorescence in a NOS-dependent manner, as responses were inhibited by L-NAME but not D-NAME (**Figure 6B**). These DAF-2 responses, likely reflecting NO diffusion into the ASL, were blocked by pretreatment (10 µM, 45 min) with GPCR G protein inhibitor YM-254890 (105–107) or HSP90 inhibitors geldanamycin, 17-AAG, or BIIB 021 (**Figure 6C**) but were not blocked by HSP70 inhibitor VER-15508. Like submerged H441s, we saw inhibition with eNOS siRNA but not with scramble, nNOS, or PAR-2 siRNA (**Figure 6D**).

**Figure 6:**
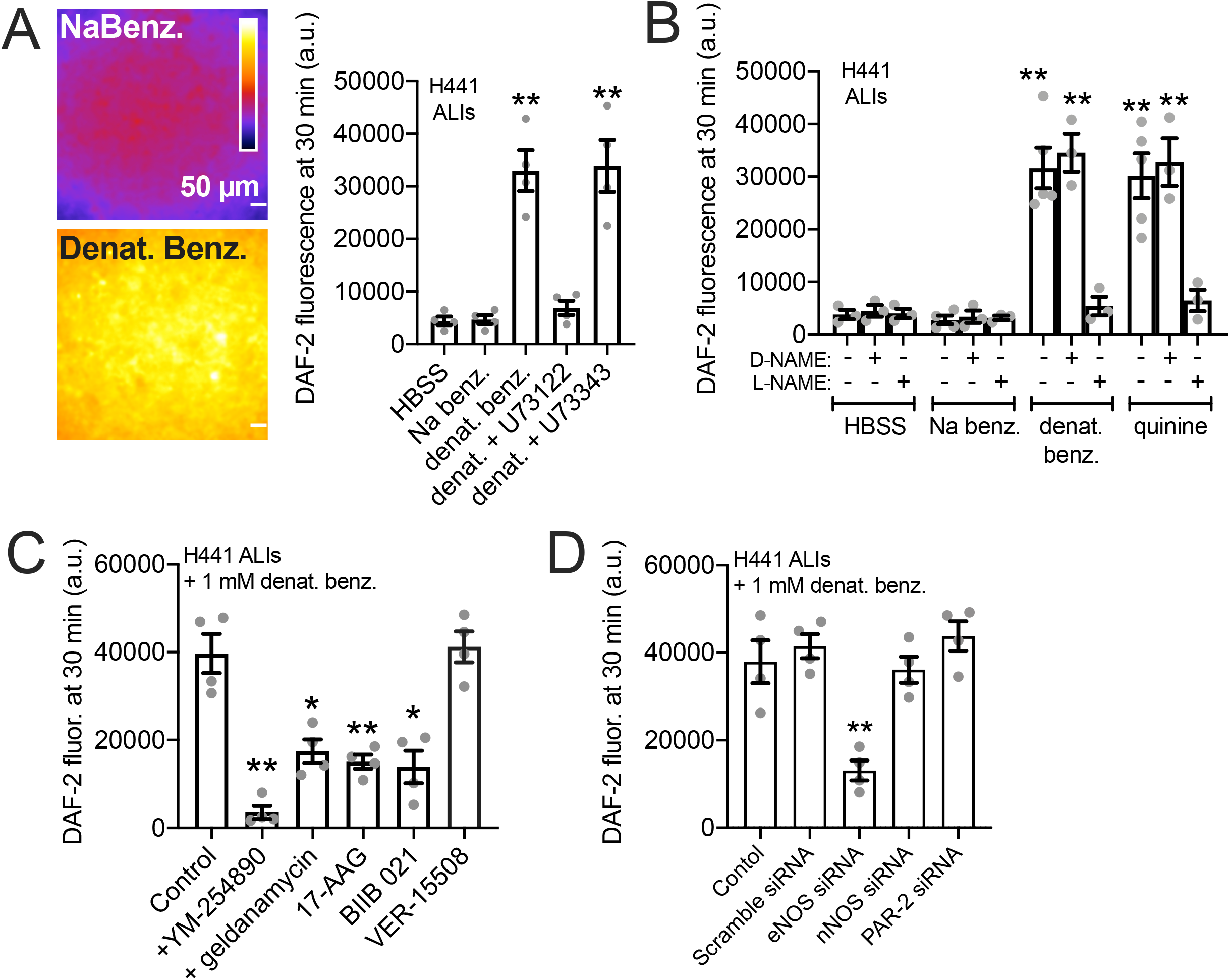
HSP90 inhibition reduces T2R stimulated NO diffusion into the airway surface liquid (ASL) in H441 cells grown at air liquid interface (ALI) ***A:*** Representative images and bar graph of 4 independent experiments of fluorescence at the apical plane of ALI when 100 µL of solution containing cell impermeable DAF-2 was placed on top (1.1 cm^2^ transwell) either containing sodium benzoate (top) or denatonium benzoate (bottom). Cultures were either pretreated with 0.1% DMSO (vehicle control), 10 µM PLC inhibitor U73122, or 10 µM inactive analogue U73343 prior to the experiment. Significance determined by one-way ANOVA with Dunnett’s posttest comparing all values to HBSS only control. ***B:*** Bar graph of experiments done as in *A* but testing inhibition of denatonium-induced or quinine-induced ASL DAF-2 fluorescence ± NOS inhibitor L-NAME or inactive D-NAME (10 µM). Bar graph is mean ± SEM of 3-5 independent experiments imaged at identical conditions. Significance by one-way ANOVA with Bonferroni posttest comparing all values to respective HBSS control; \*\**p*<0.01. ***C:*** Denatonium-stimulated H441 DAF-2 ASL fluorescence increases were reduced in the presence of GPCR signaling inhibitor YM254890 or HSP90 inhibitors geldanamycin, 17-AAG, or BIIB 021. HSP70 inhibitor VER-15508 had no effect. Bar graph is mean ± SEM of 4 independent experiments. Significance by one-way ANOVA with Dunnett’s posttest comparing all values to control (0.1% DMSO only); \**p*<0.05 and \*\**p*<0.01. ***D:*** H441s were treated with siRNA as described in the methods. ASL DAF-2 responses during denatonium stimulation were reduced by eNOS siRNA but not with scramble, nNOS, or PAR-2 siRNA. Bar graph shows mean ± SEM of 4 independent experiments (separate siRNA transfections). Significance by one-way ANOVA with Dunnett’s posttest comparing all values to control; \*\**p*<0.01.

### HSP90 inhibition reduces T2R-driven NO production in primary nasal air-liquid interface cultures

We examined NO production using DAF-FM in primary sinonasal cells grown from residual surgical material and differentiated at ALI as described (18, 19, 69). These cells express T2R receptors in apical motile cilia (**Figure 7A** and (18–21)). Note that, unlike H441 assays, denatonium benzoate was not used in primary nasal cell assays. While primary bronchial ciliated cells respond to denatonium benzoate (5), we find that nasal ciliated cells do not (18, 26), likely due to differential T2R expression between bronchial and nasal cells. For primary nasal cells, we instead used the T2R38-specific agonist phenylthiocarbamide (PTC) (70, 71) in primary ALIs genotyped for functional (PAV) or non-functional (AVI) polymorphisms in *TAS2R38* encoding the T2R38 receptor (18, 71, 108). Homozygous PAV/PAV *TAS2R38* cells produced NO in response to 1 mM PTC while AVI/AVI cells did not (**Figure 7B**). The NO produced during PTC stimulation was inhibited by geldanamycin (10 µM, 45 min pre-treatment; **Figure 7B**). We also tested the plant flavonoids apigenin and quercetin. Apigenin, a T2R14 and 39 agonist (20, 109), stimulates NO production and ciliary beat frequency increases in primary nasal ALIs via T2R14 (20). Apigenin (100 µM) stimulation increased DAF-FM fluorescence in primary nasal ALIs that was reduced by pretreatment with T2R14/39 antagonist 4’-fluoro-6-methoxyflavanone (50 µM; 45 min; (20, 110)) as well as HSP90 inhibitor geldanamycin (10 µM 45 min; **Figure 7C**). We also tested quercetin, another plant flavonoid shown to be a T2R14 agonist in heterologous expression assays (111). While quercetin was previously shown to increase (CBF) (112) and reduce cAMP signaling (113), a mechanism for these effects was not elucidated. As T2Rs also decrease cAMP through inhibitory G protein signaling in airway cells (19, 114), we hypothesized that quercetin may act as a T2R agonist in airway epithelial cells. Quercetin (50 µM) stimulation likewise increased DAF-FM fluorescence that was blocked by 4’-fluoro-6-methoxyflavanone or geldanamycin (**Figure 7D**). Apigenin and quercetin-stimulated DAF-FM fluorescence responses are summarized in **Figure 7E**.

**Figure 7.**
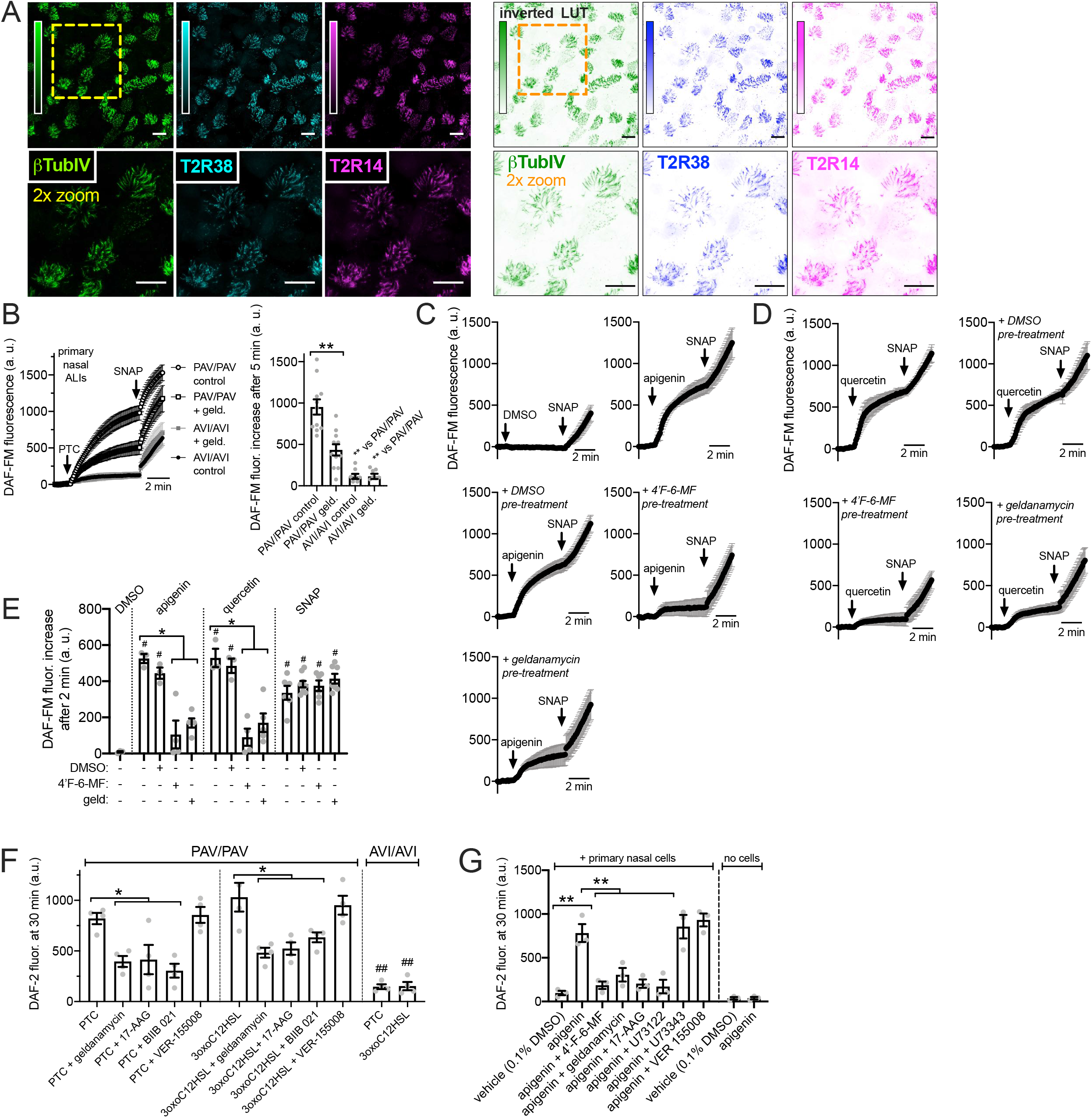
HSP90 inhibition reduces T2R-stimulated NO production in primary sinonasal epithelial cells grown at air liquid interface (ALI). ***A:*** Representative images of cilia marker β-tubulin IV (green), T2R38 (cyan) and T2R14 (magenta) immunofluorescence in an apical confocal section of primary human sinonasal ALI. Left shows conventional look-up table (LUT) and right shows inverted LUT. Scale bar is 20 µm. ***B:*** Intracellular DAF-FM increases were measured in response to T2R38-agonist PTC (1 mM) followed by NO donor SNAP (25 µM) as positive control. PTC stimulated NO production in ALIs from PAV/PAV (homozygous functional T2R38) but not AVI/AVI (homozygous nonfunctional T2R38) ALIs (nonfunctional T2R38) patients. Geldanamycin pretreatment inhibited the NO production in PAV/PAV ALIs. Trace and bar graph show mean ± SEM of 8-10 experiments per condition using ALIs from 4-5 patients. Significance determined by 1-way ANOVA with Tukey-Kramer posttest comparing all values; \*\**p*<0.01. ***C:*** Traces of DAF-FM fluorescence in PAV/AVI (heterozygous T2R38) cultures stimulated with T2R14/39 agonist apigenin (100 µM) shown with 0.1% DMSO vehicle control. Pretreatment with T2R14/39 antagonist 4’-fluoro-6-methoxyflavanone (4’-F-6-MF) or HSP90 inhibitor geldanamycin but not 0.1% DMSO (inhibitor vehicle control) reduced apigenin-induced but not SNAP-induced DAF-FM fluorescence increases. ***D:*** Traces of DAF-FM fluorescence in PAV/AVI (heterozygous T2R38) cultures stimulated with T2R14 agonist quercetin (50 µM). Pretreatment with T2R14/39 antagonist 4’-fluoro-6-methoxyflavanone (4’-F-6-MF) or HSP90 inhibitor geldanamycin but not 0.1% DMSO (inhibitor vehicle control) reduced quercetin-induced but not SNAP-induced DAF-FM fluorescence increases. ***E:*** Bar graph of intracellular DAF-FM fluorescence increases after 2 min stimulation from experiments as in *C* and *D*. Stimulation (DMSO vehicle control, apigenin, quercetin, or SNAP) listed on top and pre-treatment (DMSO vehicle control, 4’-F-6-MF, or geldanamycin) listed on the bottom. Each data point is an independent experiment (n = 4-8 per condition). Significance by Bonferroni posttest; *p<0.05 vs bracketed bars; #p<0.05 vs DMSO alone. ***F-G:*** Experiments were performed as in Fig 5 to measure NO diffusion into the ASL but with primary nasal ALIs. *F* shows PTC (500 µM) or 3oxoC12HSL (100 µM) stimulated extracellular DAF-2 fluorescence in PAV/PAV and AVI/AVI cultures, as indicated. PAV/PAV cultures were also pretreated with HSP90 inhibitors geldanamycin, 17-AAG, or BIIB 021 or HSP70 inhibitor VER-155008. *G* shows experiments with apigenin ± 4’-F-6-MF, geldanamycin, 17-AAG, or PLC inhibitor U73122 and inactive analogue U73343. Control transwells containing no cells were incubated with vehicle only or apigenin to test for any cell-independent reaction of apigenin with DAF-2.

We also performed similar assays as in H441s in Figure 5 to measure NO diffusion into the ASL using 30 µL of DAF-2 solution overlaid onto the primary nasal ALIs. Both PTC (1 mM) and *P. aeruginosa* quorum-sensing molecule 3oxoC12HSL (100 µM) increased apical surface DAF-2 fluorescence in a manner that was T2R38 dependent as it occurred in PAV/PAV (functional T2R38 homozygous) cultures but not AVI/AVI (non-functional T2R38 homozygous) cultures (**Figure 7F**). PTC- and 3oxoC12HSL-induced increases in DAF-2 fluorescence were inhibited by geldanamycin, 17-AAG, or BIIB 021 but not VER-15008 (all 10 µM for 45 min pre-treatment; **Figure 7F**). Apigenin (100 µM) increased apical DAF-2 fluorescence in a manner that was inhibited by T2R14 antagonist 4’-fluoro-6-methoxyflavanone (110) (50 µM for 45 min pre-treatment; **Figure 7G**) or PLC inhibitor U73122 (10 µM for 45 min pre-treatment; **Figure 7G**). Apigenin-stimulated DAF-2 increases were also reduced by HSP90 inhibitors geldanamycin or 17-AAG but not by HSP70 inhibitor VER 155008 (all 10 µM for 45 min pre-treatment) (**Figure 7G**). Phospholipase C (PLC) inhibitor U73122 (10 µM for 45 min pre-treatment) also inhibited the apigenin response while inactive analogue U74434 had no effect. As a control, when apigenin or vehicle was incubated in DAF-2 solution in the absence of cells (just transwells), no differences in DAF-2 fluorescence were observed (**Figure 7G**).

### HSP90 inhibition reduces T2R/NO-driven nasal ciliary beating

Data in Figure 7 suggest that HSP90 function is required for NO production during T2R38 or T2R14 activation in primary nasal epithelial cells. We tested if this affected ciliary beat frequency (CBF) using the T2R14 agonist apigenin. As previously described (20), apigenin increased ciliary beat frequency ∼10-15% over 10 min. This was blocked by T2R14 antagonist 4’-fluoro-6-methoxyfavanone or HSP90 inhibitors geldanamycin or BIIB 021 (10 minute pre-treatment, 10 µM) but not by HSP70 inhibitor VER 1555008 (**Figure 8A-B**). Thus, HSP90 inhibitors reduced apigenin-stimulated T2R14 CBF responses. We also observed a ∼30% increase in CBF with apical application of 25 µM quercetin (**Figure 8C**) that was reduced by the T2R14 inhibitor 4’-fluoro-6-methoxyflavanone (110) or geldanamycin. There was no inhibition of CBF increases in response to purinergic agonist ATP (50 µM; **Figure 8A and C**). Quercetin-stimulated increases in CBF were also inhibited by blocking NO signaling with L-NAME (10 µM; **Figure 8D**). These data suggest quercetin activation of CBF may occur through T2R activation and NO production. Importantly, we observed that geldanamycin has no significant effect on baseline CBF after ≥20 min (**Figure 8E**), in contrast to prior studies in mouse tracheal cells, where geldanamycin rapidly reduced CBF to ∼75% of basal values, postulated to be due to reduced stability of tubulin polymerization upon HSP90 inhibition (65). We did not see these effects.

**Figure 8.**
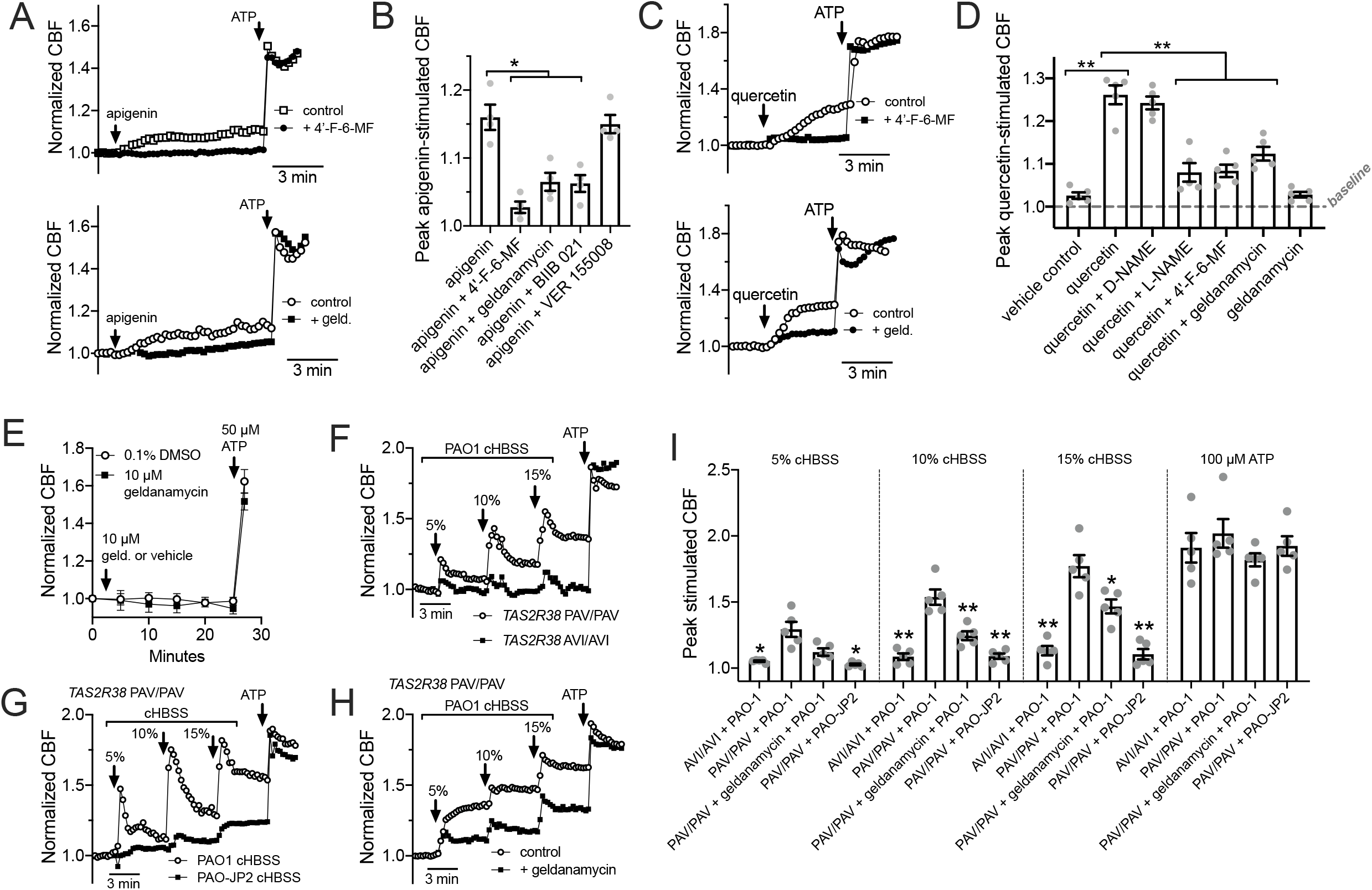
HSP90 inhibition reduces T2R-stimulated ciliary beating in primary sinonasal epithelial cells. **A:** Left shows representative normalized CBF responses (representative experiments shown) to T2R14/39 agonist apigenin in human primary sinonasal ALIs ± T2R14/39 inhibitor 4’-fluoro-6-methoxyflavanone. Right shows normalized CBF responses (representative experiments shown) to apigenin ± geldanamycin (10 µM; 5 min pre-treatment). Mean baseline CBF was not with vehicle or 4’-fluoro-6-methoxyflavanone pre-treatment (7.5 ± 1.1 Hz or 8.2 ± 0.9 Hz, respectively; not significant by Students’ *t* test). Mean baseline CBF was also not different before or after vehicle or geldanamycin pre-treatment (6.9 ± 1.7 Hz or 7.9 ± 1.2 Hz; respectively; not significant by Students’ *t* test). ***B:*** Bar graph of mean ± SEM of CBF responses from 5 independent experiments as shown in *A* using ALIs from 4 different patients. Significance determined by 1-way ANOVA with Bonferroni posttest; \**p*<0.05. ***C:*** Left shows representative normalized CBF responses (representative experiments shown) to T2R14/39 agonist quercetin in human ALIs ± T2R14/39 inhibitor 4’-fluoro-6-methoxyflavanone. Mean baseline CBF was not with vehicle or 4’-fluoro-6-methoxyflavanone pre-treatment (7.3 ± 1.2 Hz or 7.9 ± 0.6 Hz, respectively; not significant by Students’ *t* test). Right shows normalized CBF responses (representative experiments shown) to quercetin ± geldanamycin (10 µM; 5 min pre-treatment). Mean baseline CBF was not different before or after vehicle or geldanamycin pre-treatment (7.4 ± 1.3 Hz or 7.0 ± 0.9 Hz; respectively; not significant by Students’ *t* test). ***D:*** Bar graph of mean ± SEM of CBF responses from 5 independent experiments as shown in *C* using ALIs from 5 different patients. Significance determined by 1-way ANOVA with Bonferroni posttest; \*\**p*<0.01. ***E:*** Graph shows real-time measurement of CBF (mean ± SEM of 6 independent experiments using ALIs from 3 patients) during prolonged geldanamycin treatment, followed by stimulation with purinergic agonist ATP. ***F*:** Primary nasal ALIs genotyped for functional T2R38 (TAS2R38 PAV/PAV) or non-functional T2R38 (TAS2R38 AVI/AVI) were stimulated with diluted HBSS in which P. aeruginosa PAO-1 had been incubated overnight (conditioned HBSS; cHBSS, diluted with unconditioned HBSS). Peak CBF responses to PAO-1 cHBSS were greater in PAV/PAV cells vs AVI/AVI cells. Representative trace shown from 5 experiments using cultures from separate individual patients. ***G*:** PAV/PAV cells were stimulated with cHBSS from PAO-1 or PAO-JP2, which lacks the ability to produce AHLs. PAO-1 cHBSS stimulated CBF increases that were greater than CBF increases observed with PAO-JP2 cHBSS. Representative trace shown from 5 experiments using cultures from separate individual patients. ***H***: PAV/PAV cells were stimulated with PAO-1 cHBSS ± geldanamycin pretreatment. Representative trace shown from 5 experiments using cultures from separate individual patients. ***I***: Bar graph showing peak CBF (mean ± SEM with individual data points showing individual experiments) observed from experiments as in *F-H*. Asterisks represent significance compared with PAV/PAV + PAO-1 cHBSS at each individual concentration, determined by Sidak’s multiple comparison test; \**p*<0.05 and \*\**p*<0.01.

We also tested CBF response to HBSS that had been conditioned by overnight exposure to P. aeruginosa. We previously performed similar experiments with conditioned LB media and showed that CBF increases in response to dilute (6.25%-12%) P. aeruginosa media were dependent on bitter receptor T2R38, which is expressed in cilia and detects acylhomoserine lactone (AHL) quorum sensing molecules (18). Here, *P. aeruginosa* Wt strain PAO-1 was incubated in HBSS for 24 hrs, and the resulting conditioned HBSS (cHBSS) was diluted and used to stimulated cells. We found that 5-15% cHBSS stimulated robust ciliary responses in nasal ALIs homozygous for the functional polymorphism (PAV) of the TAS2R38 gene encoding the T2R38 receptor (**Figure 8F**). Cells homozygous for the non-functional (AVI) polymorphism of TAS2R38 responded with much lower CBF increases (**Figure 8F**), showing the responses were dependent on T2R38. With cHBSS from strain PAO-JP2, which is unable to produce AHLs (78, 79, 115), we observed minimal CBF responses in PAV/PAV cells compared with PAO-1 Wt cHBSS (**Figure 8G**), showing the response were dependent on AHL signaling. Notably, AHL signaling also control production of quinolone quorum sensing molecules (116), which can also function as T2R agonists (19). Fitting with a role for HSP90 in T2R function, we observed that geldanamycin reduced the CBF response to PAO-1 cHBSS in PAV/PAV cells (**Figure 8H**). These data are summarized in **Figure 8I** and together suggest that geldanamycin can reduce the ability of nasal ALIs to detect *P. aeruginosa* through T2Rs and increase ciliary beating.

### HSP90 inhibition reduces T2R NO production and phagocytosis in primary human macrophages

We wanted to examine T2R signaling to eNOS in another human primary cell model to test if it requires HSP90 function. Like epithelial cells, macrophages are important players in early innate immunity. Unprimed (M0) monocyte-derived macrophages also express eNOS involved in enhancement of phagocytosis during immune receptor activation (33). While isolated monocytes differentiate into Mϕs that are not exactly the same as alveolar Mϕs that populate the airways at baseline (117–119), monocyte-derived Mϕs are often used as surrogates for alveolar Mϕs and are nonetheless themselves important for infections, including during chronic airway inflammation like CRS, chronic obstructive pulmonary disease (COPD), asthma, and cystic fibrosis (17, 120, 121). We previously observed that T2R stimulation in human M0 monocyte-derived macrophages also activates low-level Ca^2+^ responses that drive NO production to enhance phagocytosis (68). Macrophage DAF-FM responses to denatonium benzoate were significantly reduced by HSP90 inhibitors geldanamycin and BIIB 021 (10 µM; 30 min pre-treatment; **Figure 9A**) despite no change in denatonium-induced Ca^2+^ signals (**Figure 9B**), suggesting that HSP90 is required for activation of eNOS and/or nNOS downstream of the T2R-induced Ca^2+^ response (68).

**Figure 9.**
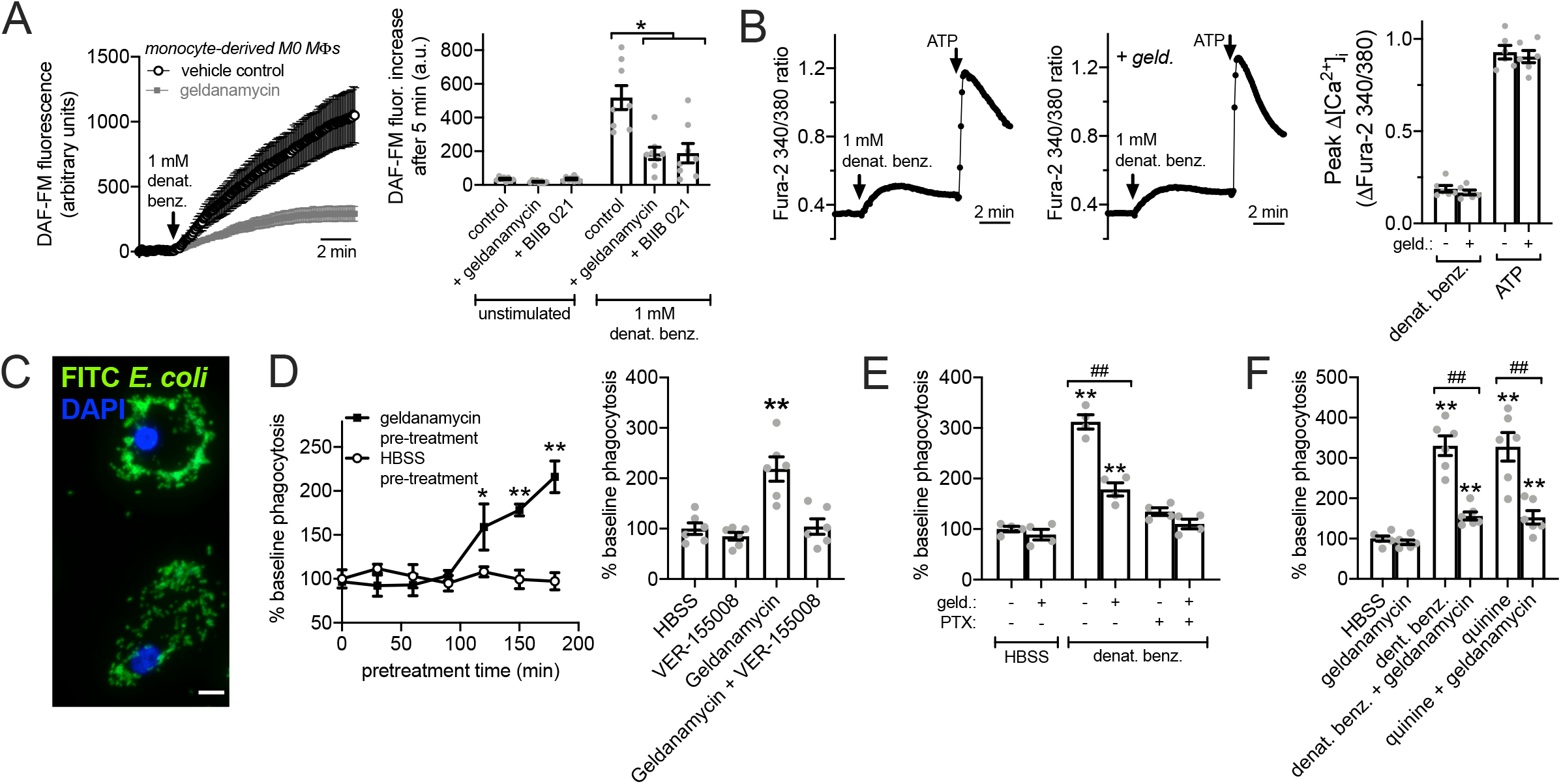
HSP90 inhibition reduces T2R-stimulated NO production and FITC-*E. coli* phagocytosis in primary human M0 macrophages. ***A*:** DAF-FM-loaded macrophages exhibited increases in fluorescence in response to 1 mM denatonium benzoate that were strongly inhibited by geldanamycin. Left shows average traces and right shows bar graphs (mean ± SEM) from 8 independent experiments using macrophages from two donors. DAF-FM fluorescence increase was also inhibited by BIIB 021. Control = denatonium benzoate after pretreatment with 0.1% DMSO. ***B:*** Low-level Ca^2+^ responses to denatonium benzoate were not affected by geldanamycin. Left shows representative traces in the absence or presence of 1 µM geldanamycin. Right shows bar graph of 6 independent experiments using macrophages from three different donors. Response to purinergic agonist ATP shown as control. ***C:*** Representative image of macrophages with phagocytosed FITC-labeled *E. coli*. ***D:*** Left shows time course of phagocytosis responses during 30 min incubation in HBSS as described in the methods after pre-treatment with geldanamycin or HBSS for times indicated on the y-axis. Each data point is mean ± SEM of 3 independent experiments using macrophages from 3 different donors. Right shows separate experiments of baseline phagocytosis over 30 min (HBSS only) of FITC-*E. coli* after 2 hours pre-treatment with HBSS only (containing 0.1% DMSO as vehicle control), 1 µM VER-15508, 1 µM geldanamycin, or geldanamycin plus VER-15508. Significance determined by 1-way ANOVA with Dunnett’s posttest comparing values to HBSS pretreatment; **p <0.01. Bar graph shows mean ± SEM of 6 experiments using macrophages from 3 donors. ***E:*** Stimulated 30 min phagocytosis of FITC-*E. coli* (HBSS only control or 1 mM denat. benz. ± pertussis toxin [PTX]) was measured after pre-incubation with HBSS + 0.1% DMSO or 1 µM geldanamycin. PTX and geldanamycin both inhibited denatonium-induced phagocytosis. Significance determined by 1-way ANOVA with Bonferroni posttest; \*\**p*<0.01 vs HBSS control and ^##^*p*<0.01 vs bracketed groups. ***F:*** Geldanamycin reduced phagocytosis increases observed with both denatonium and quinine. Bar graph shows mean ± SEM of 6 independent experiments using cells from 6 different individual patients. Significance by one way ANOVA with Tukey-Kramer posttest comparing all bars; \*\**p*<0.01 vs HBSS alone; ^##^*p*<0.01 vs bracketed bar.

We measured phagocytosis of FITC-labeled *Escherichia coli* (**Figure 9C**). First, we had to test effects of geldanamycin alone. It was previously shown that geldanamycin treatment and HSP90 inhibition increases phagocytosis after ∼90 minutes due to transcriptional up-regulation of HSP70 (122, 123). We observed an increase in baseline phagocytosis after ∼2 hours geldanamycin treatment (**Figure 9D**); this was inhibited by HSP70 inhibitor VER-155008 (**Figure 9D**), supporting these prior observations. Thus, to avoid any effects of HSP70 up-regulation, we tested the effects of geldanamycin on denatonium-upregulated phagocytosis after only 30 min geldanamycin pretreatment followed by continued geldanamycin treatment for the 15 minutes of the phagocytosis assay (45 min total). We observed a ∼3 fold increase in phagocytosis in response to 1 mM denatonium benzoate (as we previously reported (42)) that was inhibited by geldanamycin as well as pertussis toxin (**Figure 9E**), which inactivates the G^i^ and G^gustducin^ Gα subunits that can couple to T2R receptors (114, 124, 125). We also saw inhibition of denatonium benzoate-induced or quinine (500 µM)-induced phagocytosis of pHrodo-labeled *S. aureus* with geldanamycin pretreatment (**Figure 9F**). Together, these data suggest that HSP90 plays a key role in activation of NO production downstream of T2R activation.

To confirm that our FITC-*E. coli* measurements reflected phagocytosis and to test a pathogen with more relevance to the airway epithelium, we also tested pHrodo-labeled *Staphylococcus aureus*, as previously utilized in (42). The pHrodo dye fluorescence strongly in acidic environments and thus exhibits a marked increase in fluorescence when internalized into acidic organelles like lysosomes and phagosomes. Assays were carried out similarly to FITC-*E. coli* assays described above. We observed that denatonium benzoate (1 mM) increased phagocytosis in a NOS-dependent manner as it was inhibited by L-NAME but not D-NAME (10 µM, 30 min pre-treatment; **Figure 10A-B**). The increased phagocytosis in response to denatonium benzoate or *P. aeruginosa* 3oxoC12HSL (100 µM) was inhibited by geldanamycin or BIIB 021 (10 µM pretreatment; **Figure 10C-D**), supporting reduction of this innate immune response by HSP90 inhibitors. We tested other HSP90 inhibitors using pHrodo *S. aureus* in the same assay in a plate reader format (as described in the methods and (42)). Well fluorescence increased when macrophages were incubated with 1 mM denatonium benzoate for 15 min (**Figure 10E**). Equimolar sodium benzoate had no effect (**Figure 10E**). The stimulatory effect of denatonium benzoate was reduced by pertussis toxin (to block T2R GPCR signaling) or pre-incubation (15 min; 10 µM) with geldanamycin, BIIB 021, or 17-AAG but not VER 155008 (**Figure 10E**).

**Figure 10.**
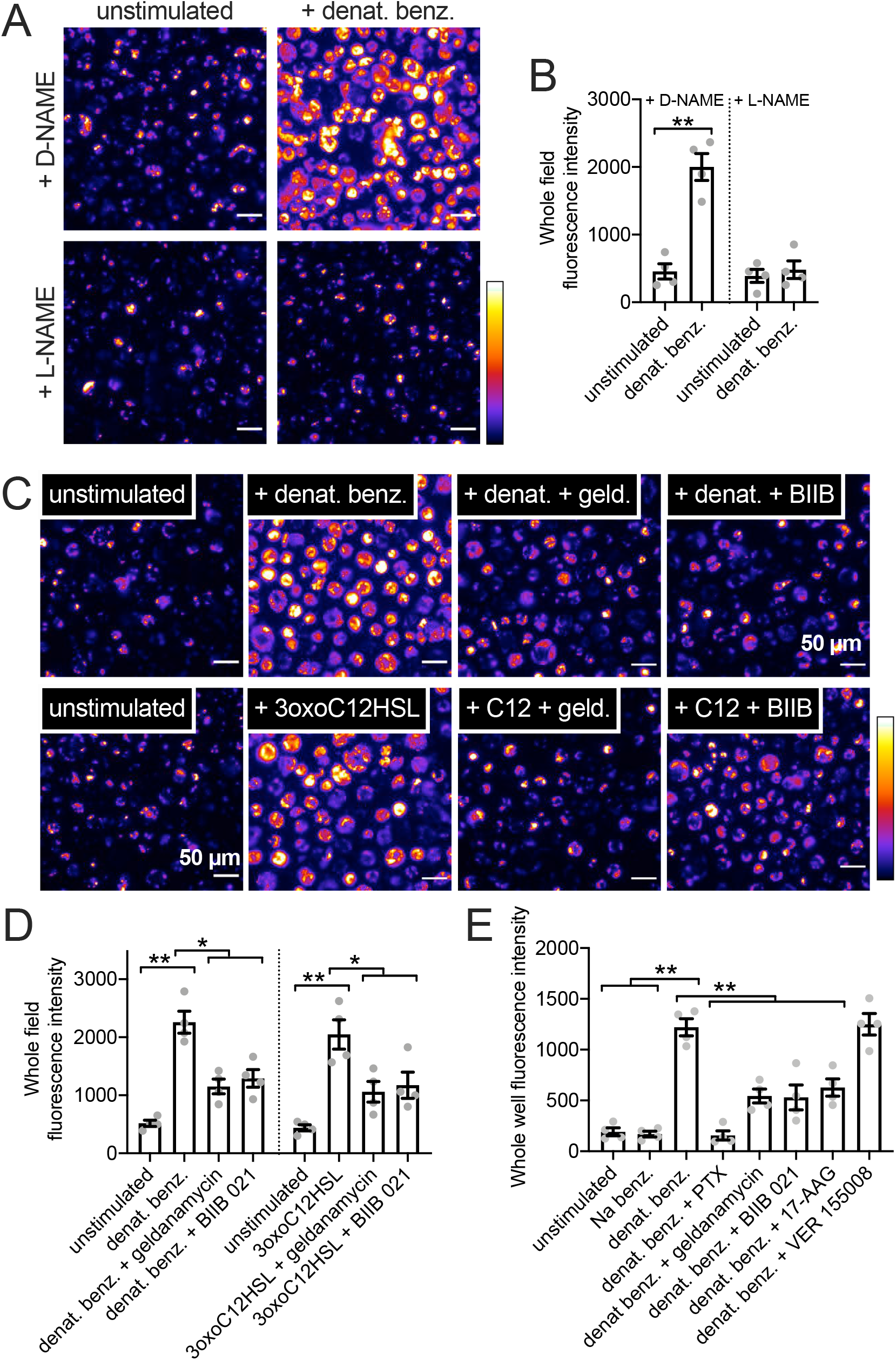
HSP90 inhibition reduces T2R-stimulated pHrodo-*S. aureus* phagocytic responses in primary human M0 macrophages. ***A:*** Representative images of pHrodo-labeled *S. aureus* phagocytosis in primary human macrophages ± denatonium benzoate (1 mM) stimulation after D-NAME or L-NAME pre-treatment (10 µM; 45 min). ***B:*** Bar graph of pHrodo-*S. aureus* fluorescence after experiments as in *A*. Significance by Bonferroni posttest with paired comparisons; \*\**p*<0.01. ***C:*** Representative images of pHrodo-labeled *S. aureus* phagocytosis in primary human macrophages ± denatonium benzoate (1 mM) or 3oxoC12HSL (100 µM) after no-pretreatment (0.1% DMSO only as vehicle control)) or pretreatment with HSP90 inhibitors geldanamycin or BIIB 021 (pretreatment as in Fig 8). ***D:*** Bar graph of pHrodo-*S. aureus* phagocytosis during stimulation with HBSS only (unstimulated control), 1 mM denatonium benzoate, or 100 µM 3oxoC12HSL ± geldanamycin or BIIB 021 (pretreatment as in Fig 9). Significance by one-way ANOVA with Bonferroni posttest; \**p*<0.05 or \*\**p*<0.01. ***E:*** Bar graph of pHrodo-*S. aureus* phagocytosis during stimulation with HBSS only (unstimulated control) or 1 mM denatonium benzoate ± pertussis toxin (PTX), geldanamycin, BIIB 021, 17-AAG, or VER 15508. PTX (500 ng/ml) pre-treatment was 18 hrs. Macrophages were pre-treated with other inhibitors as in Fig 9. Significance by one-way ANOVA with Bonferroni posttest; \**p*<0.05 or \*\**p*<0.01.

## DISCUSSION

HSP90 likely plays a multi-faceted role in airway epithelial physiology beyond facilitating protein folding, but data surrounding its specific contributions are unclear. There has been an increasing interest in heat shock chaperone proteins in immune cell modulation (126), including regulation of immune cell metabolism (127) and immune receptor signaling (128, 129). Here, we show a new innate immune role for HSP90, namely the production of NO downstream of T2R signaling. Specifically, we show that HSP90 inhibition by multiple structurally diverse compounds acutely impairs NO-mediated airway epithelial CBF responses and macrophage phagocytosis without impairing the upstream calcium signaling. Thus, the result of HSP90 inhibition is not simply impaired receptor function, trafficking, or folding. We utilized several models, from heterologous expression in HEK293T and A549 cells to human primary cells differentiated from patient material.

The T2R to eNOS pathway, specifically polymorphisms regulating T2R38 signaling, has been identified as clinically important in terms of increased susceptibility to upper respiratory infections and impaired patient outcomes in chronic rhinosinusitis (CRS) (45-48, 50-52, 130-132). T2R signaling to eNOS regulates both airway ciliary beating and macrophage phagocytosis. Others have shown that HSP90 is important for scaffolding eNOS with activating proteins like Akt or Ca^2+^-bound calmodulin. We hypothesize that this scaffolding function is likewise important for T2R activation of eNOS. This puts HSP90 in a prime role to regulate eNOS output during T2R stimulation. HSP90 transcript levels can be regulated by a host of transcription factors active during inflammation or cell stress, including HSF1, NF-IL6, and NFκB (133). Various posttranscriptional modifications, from phosphorylation to nitrosylation (133, 134), can also alter HSP90 activity. We hypothesize that regulation of HSP90 may be one way to modulate airway T2R/eNOS NO output or airway NO output in general. In a myocardial ischemia-reperfusion injury mouse model, transfection of HSP90 is protective by enhancing eNOS S1177 (activating site) phosphorylation and decreasing eNOS T495 (inhibitory site) phosphorylation (135). We hypothesize that HSP90 expression might be a pathway that could be exploited to increase NO in airway diseases associated with reduced NO levels, including CF (136–144) or primary ciliary dyskinesia (145–147).

While HSP90 has been shown to be important for baseline cilia function (63–65), other studies have suggested it is necessary for Th2 (IL-13-driven) and Th17 (IL-17-driven) airway goblet cell metaplasia (66). Many airway diseases, including asthma, COPD, and CF, are characterized by a loss of ciliated cells due to goblet or squamous metaplasia, likely impairing mucociliary clearance both through mucin hypersecretion and loss of cilia (148). HSP90 inhibition was suggested to be potentially useful in type 2 inflammatory airway disease characterized by airway remodeling, typically goblet cell metaplasia (66), which include asthma and chronic rhinosinusitis. However, outside the airway, HSP90 has been implicated in both pro-inflammatory and anti-inflammatory processes (126). A better understanding is needed of how HSP90 contributes to the myriad of functions that airway epithelial cells perform, including bacterial surveillance and antimicrobial responses.

While HSP90 is required for NO production during stimulation of airway and macrophage T2R s, the inhibition of HSP90 did not affect T2R-mediated Ca^2+^ signals upstream of the NO production. HSP90 inhibition did reduce cGMP production, elevation of ciliary beating, and phagocytosis, all downstream of NO production. Thus, HSP90 inhibition may reduce innate immune responses to bacteria in the airway, through both epithelial cells and dedicated immune cells. The overall effect on airway innate immunity will depend on other pathways that are up or down regulated. It may be that the reversal of goblet metaplasia in asthma with HSP90 inhibition outweighs a side-effect of reduced T2R responses, but the knowledge that these T2R responses are reduced may suggest other supplemental targets/therapies are needed to boost NO production in patients receiving HSP90 inhibitors. Here, we simply show one effect of HSP90 inhibition that would be predicted to be detrimental. Other effects of HSP90 inhibitors in the airway must be studied in more detail to clarify the entire picture of how these drugs may affect the respiratory epithelium and innate defense.

As described above, HSP90 has been localized to the base of airway cilia (63, 64) in close proximity to eNOS (29, 32, 64, 149) and T2Rs (5, 18–20). We found that T2R activation produces more NO in ciliated airway cells than stimulation of purinergic receptors (18) or PAR-2 (69), despite the fact that these other GPCRs generate higher Ca^2+^ responses than T2Rs. A clearer picture of the T2R signaling pathway is necessary to understand why this occurs. It may be that the close proximity of T2Rs to eNOS and HSP90 creates localized Ca^2+^ or calmodulin microdomains within the cilia. Another explanation is that other T2R-stimulated pathways downstream or in parallel to the Ca^2+^ also contribute to these responses. While eNOS can be activated directly through interactions with Ca^2+^-bound calmodulin (150, 151), it can also be activated by phosphorylation at multiple residues by kinases like Akt, CaMKII, or PKA (152–161). Phosphorylation at one or more eNOS residues may be important during T2R activation of NO production. While much work on T2Rs has focused on Ca^2+^ activation downstream of the Gβγ component of their heterotrimeric G protein signal pathway, little is known about kinases activated during T2R stimulation. Future studies are needed to better elucidate the molecular mechanisms of T2R signaling to eNOS in airway epithelial cells and other cells like macrophages.

## Acknowledgements

We thank M. Victoria (University of Pennsylvania) for excellent technical assistance.

## Disclaimers

The content is solely the responsibility of the authors and does not necessarily represent the official views of the National Institutes of Health.

## Author contributions

R.M.C., B.M.H., and R.J.L. performed experiments and analyzed data. R.M.C. and R.J.L. wrote the paper. N.D.A., J.N.P. recruited patients, aided with tissue procurement and regulatory approvals, facilitated culture of primary human cells, maintained clinical databases and records, and intellectually contributed. All authors reviewed and approved the final manuscript.

